# Influence of high fibre diets on the gut microbiota, prostate tumour growth and normal tissue toxicity following ionising radiation

**DOI:** 10.64898/2026.08.13.744579

**Authors:** Aliu Moomin, Carlos Sabater, Merel A van den Haak, Adam Potter, Susan M Hay, David McClelland, Elaina SR Collie Duguid, Heather M Wilson, Anne E Kiltie

## Abstract

**Purpose:** High dietary fibre intake has been linked to lower cancer risk, yet its role in prostate cancer treatment responses and radiotherapy tolerance remains unclear. We evaluated the effects of dietary fibres (inulin, pectin, β-glucan) on prostate tumour growth, gut microbiota and intestinal response to ionising radiation (IR) in murine models.

**Methods:** Male FVB and C57BL/6J mice were injected with murine Myc-CaP (FVB), RM-1 or DVL3 (C57BL/6J) prostate tumour cells and fed a low-fibre (0.2% cellulose) or high-fibre diet (10% inulin, pectin or β-glucan). Some mice had tumour irradiation (6 Gy). Tumour volume, caecal weight and faecal microbiota relative abundance (by 16S rRNA gene sequencing) were analysed. Caecal contents fermentation acids were quantified by gas chromatography. The effects of dietary fibre on intestinal acute normal tissue toxicity post-irradiation (10–14 Gy) were assessed by intestinal crypt assay.

**Results:** Inulin delayed average tumour growth in all models. Inulin and β-glucan prolonged post-IR tumour control versus 0.2% cellulose (all p <0.05), in some but not all mice. Inulin, pectin and β-glucan increased faecal acetate concentrations post-IR and mice demonstrated responder (R) vs non-responder (NR) phenotypes to diet/IR, associated with *Bifidobacterium* (inulin-R), *Lactobacillus* and *Parasutterella* (pectin-R) and *Muribaculacaeae* and *Muribaculum* (β-glucan-R). High fibre-fed mice had enhanced intestinal crypt regeneration following 12 Gy compared to 0.2% cellulose-fed mice.

**Conclusions:** High fibre diets slowed prostate tumour growth both alone and following 6 Gy IR, while protecting small intestines from radiation-induced injury. Effects may have been mediated via increased microbiota-driven metabolite production and enhanced epithelial regeneration, but more mechanistic work is required to explore causality. The differences in individual responses to various fibres should be investigated further, as this may have relevance to adopting dietary fibre supplementation strategies in human radiotherapy patients, and may reflect the recognised importance of an individual’s baseline microbiota on dietary effects.

## Introduction

Prostate cancer is the most prevalent male cancer and is the fifth leading cause of cancer deaths worldwide (1). In 2018, over 1.2 million new cases of prostate cancer were diagnosed worldwide, with more than 360,000 deaths (1,2). Between 2021 and 2022, the UK reported over 50,000 newly diagnosed cases of prostate cancer, representing a 27% increase in incidence compared to 2019 and causing over 16,000 deaths (3). This sharp rise in prostate cancer prevalence and incidence among men is partly due to increased diagnosis through prostate-specific antigen (PSA) testing and increased life expectancy (4,5).

Western lifestyles can be characterised by a positive energy balance, low physical activity, and a diet low in dietary fibre but high in refined sugars, animal protein and saturated fats (5). The UK National Diet and Nutrition Survey for 2019–2023 reported fibre intake in the UK fell below the government’s recommended levels (30 g/day) for all sex groups, with only 4% of adults (19 years and above) meeting the recommended intake of fibre (6,7).

There is growing interest in the association between dietary fibre and human health. Epidemiological and intervention studies have reported the impact of dietary fibres on prostate cancer growth. In a US population-based study, consuming more fibre was linked to a lower risk of aggressive prostate cancer, with odds ratios 0.70 (95% CI 0.50-0.97) and 0.61 (95% CI 0.40-0.93) for the second and third tertiles, respectively (8). In an intervention study with modified citrus pectin, 78% of men with biochemically relapsed, non-metastatic prostate cancer demonstrated a treatment response: 58% showed decreased PSA and 75% showed improvement in PSA doubling time (p=0.003) (9). More recently, Thomas *et al*, randomised men on active surveillance with rising PSA values to phytochemical-rich food capsules (six foods) plus or minus an additional twice daily capsule containing five probiotics, vitamin D3 and 100 mg inulin, thus slowing PSA progression rates, but whether the inulin fibre itself had an impact is unclear (10).

Furthermore, pre-clinical studies suggest that specific fibre types may also modulate tumour responses to radiotherapy. In a murine urological cancer model, we fed psyllium, psyllium plus resistant starch, or psyllium plus inulin to female C57BL/6J mice bearing UPPL1591 mouse bladder tumours. We observed psyllium plus resistant starch acted as a radiosensitiser whereas psyllium plus inulin delayed tumour growth following ionising radiation (IR) (11). Approximately 80% of patients receiving pelvic radiotherapy will develop acute radiation enteritis (radiotherapy induced damage to the intestinal epithelium), negatively affecting their quality of life (12). In mice also, dietary fibres can alleviate damage to the GI tract following exposure to ionising radiation, with mice fed a high fibre diet showing more regenerating intestinal crypts 3.75 days post-irradiation than mice fed a low fibre diet (11).

Having previously undertaken work in mouse bladder cancer flank allografts (11,13) we wished to identify a mouse prostate cancer allograft model suitable for dietary fibre and radiation studies. This is due to the large number of men with prostate cancer who could potentially benefit from taking dietary fibre supplements. For men on active surveillance, fibre supplements could slow tumour progression and delay time to prostatectomy or radiotherapy; in men having radiotherapy, it could reduce intestinal side effects and improve tumour responses. Mouse models allow more rapid, detailed investigations of the effects of fibre compared to human studies, where randomised controlled trials need large numbers of participants recruited over many months or years to be sufficiently powered.

We hypothesised that dietary fibre supplementation (with inulin, pectin and β-glucan), alone and in combination with ionising radiation (IR), would slow the growth of mouse prostate cancer flank syngeneic tumours and alleviate damage to the small intestine following exposure to IR, with alterations to the gut microbiota community and resulting metabolite production.

## Methods and Materials

### Cell culture

The murine prostate cancer cell lines Myc-CaP and RM-1 were purchased from American Type Culture Collection (ATCC, USA), and DVL3 cells were a generous donation from Professor Ian G Mills (University of Oxford, UK). All cell lines were grown in high glucose, pyruvate-supplemented Dulbecco’s modified Eagle medium (Thermofisher, UK), supplemented with 10% foetal bovine serum (Invitrogen), and cultured in a humidified atmosphere of 5% CO_2_ at 37°C. All cells tested negative for mycoplasma.

### Animals

All animal work was performed following UK Home Office Guidelines and approved by the University of Aberdeen Animal Welfare and Ethical Review Body under project licences P8484EDAE and PP9835550. FVB and C57BL/6J mice were purchased from Charles Rivers Laboratories (UK) and housed in individually ventilated cages (IVCs). All mice were acclimatised for a week and housed in a temperature-controlled environment with a 12-h reversed-phase light/dark cycle (lights on 07:00 h) and provided with food and water *ad libitum*. Mouse cages were randomised into treatment groups using the rand() function in Microsoft Excel.

### Tumour growth with modified diets

Tumours were induced by injecting DVL3 (1×10^6^) or RM-1 (0.5×10^6^) cells subcutaneously into the flanks of 6-week-old C57BL/6J mice, or Myc-CaP (1×10^6^) cells subcutaneously into the flanks of 6-week-old FVB male mice, as described in (Then et al 2024) (11). Mouse diets were changed from normal chow to modified diets (Research Diets Inc., USA), either low-fibre diet (0.2% cellulose) or high-fibre (10% inulin), see Table S1 for diet formulations. Due to the fast-growing nature of RM-1 cells, diets were changed two weeks before tumour induction, while diets were changed on the same day of tumour induction for Myc-CaP and DVL3 cells to accommodate their different growth rates. DVL3 tumours grew more slowly than anticipated and diet was changed back to normal chow 34 days after tumour injection. After tumour establishment, tumour growth was measured three times per week (daily for RM-1 model) using callipers and calculated using the formula (length x width x height) x (π/6) (11). Mice were euthanised when the tumours reached approximately 800 mm^3^.

### Tumour growth with modified diets and irradiation

Tumours were induced by injecting 1×10^6^ Myc-CaP cells subcutaneously into the flanks of 6-week-old FVB male mice, fed 0.2% cellulose or 10% inulin as described above. RM-1 (0.5×10^6^) cells were injected subcutaneously into 6-week-old C57BL/6J male mice two weeks after the mouse diets were changed from normal chow to modified diets (Research Diets Inc., USA): low-fibre diet (0.2% cellulose) or high-fibre (10% inulin, 10% apple pectin (Sigma) or 10% β-glucan (Orafti®β-fit, generous gift from Beneo)); diet formulations Table S1. After tumour establishment, tumour growth was measured three times per week (daily for RM-1 model) and mouse tumours were irradiated to 6 Gy using an Xstrahl X-ray irradiator at 300 kV and 10 mA with 1.0 mm Cu filter and 1.5 cm applicator (CIX3, Xstrahl, UK) when tumours reached 80 – 100 mm^3^. Mice were anaesthetised and blood collected by cardiac puncture followed by cervical dislocation when the tumours reached approximately 800 mm^3^ (11). Caeca were collected and weighed. Mouse faecal samples were collected for 16S rRNA gene sequencing and caecal contents for fermentation acid analysis. Faecal samples and caecal contents were transported on ice and stored at -70°C for analysis.

### DNA extraction for 16S rRNA gene sequencing

DNA was extracted from mouse faecal samples using DNeasy PowerSoil Pro DNA isolation kits according to manufacturer’s instructions (Qiagen Ltd, Manchester, UK). DNA concentration was quantified using Qubit (Qubit dsDNA BR assay kit, ThermoFisher Scientific, UK). Bacterial communities from the samples were profiled using 16S rRNA gene sequencing of the V1-V2 region on a MiSeq platform (Illumina, Inc., San Diego, CA) with v3 chemistry and 300bp paired-end reads, at the Centre for Genome Enabled Biology and Medicine, University of Aberdeen. Region-specific primers (11) which included partial Illumina adapters, were utilised to amplify the V1-V2 region. This was followed by 8 cycle PCR to add full-length Illumina adapters and dual barcodes. Equimolar quantities of the resulting libraries were pooled and sequenced on a MiSeq platform (11).

### Fermentation acid analysis

Mouse caecal contents were processed for fermentation acid (including short chain fatty acid (SCFA), lactate and succinate) analysis as described by Richardson et al., 1989 (14); 2-ethyl butyric acid (0.1 M) was used as the internal standard, with N-tert-butyldimethylsilyl-N-methyltrifluoroacetamide (MTBSTFA, Merck, UK) as derivatising agent. The caecal content from each mouse was vortexed in 2 ml PBS (supplemented with 30% glycerol) and centrifuged at 3000 rpm for 5 minutes. Concentrated HCl (0.5 ml) and ethyl ether (2 ml) were added to the supernatant (1 ml), vortexed for 1 min and centrifuged at 3000 rpm for 10 minutes. The ether layer was transferred into a 15 ml tube. The extraction of the caecal supernatant with 2 ml ethyl ether was repeated and the ether extracts were pooled. The ether extract (800 µl) was transferred into a Wheaton vial containing 100 µl of MTBSTFA, screw capped, heated on a heating block at 80°C for 20 minutes and left at room temperature for 48 hours to derivatise (11). SCFA concentrations were measured using gas chromatography on an Agilent HP 6890 (Agilent Technologies; Alpharetta, USA). Simple linear regression was used to determine the association between specific SCFA concentrations and tumour growth.

### Normal tissue response to ionising radiation

C57BL/6J mice were fed with modified diets: low-fibre diet (0.2% cellulose) or high-fibre (10% inulin, 10% pectin or 10% β-glucan) for 2 weeks. Mice were irradiated supine at 10, 12 and 14 Gy to the lower abdomen using an Xstrahl X-ray irradiator at 300 kV and 10 mA with 1.0 mm Cu filter and 3.5 cm applicator (CIX3, Xstrahl, UK). Mice were killed at 3.75 days and the intestines collected. Swiss rolls were made from the small intestines, and formalin-fixed, before being paraffin embedded and haematoxylin and eosin stained for assessment of intestinal crypt regeneration (15,16). For each sample, regenerating crypts were counted in a total of 6 mm length of damage, using QuPath software version 0.6 for analysis (17), and the percentage of regenerating crypts was calculated compared to the mean number of crypts per 6 mm length in four unirradiated controls fed the same diet.

### Data analysis and bioinformatics

General statistical analyses were performed using GraphPad Prism software (version 11.0.2) unless otherwise specified. QuPath image analysis software (version 0.6) was used for histopathological analysis of acute toxicity of ionising radiation in normal intestinal tissues from mice (17). Details are provided in the relevant figure legends and text.

Amplicon sequencing data were analysed using QIIME2 v.2021.8 software (18). Raw paired-end sequences were first quality filtered, retaining sequences with a mean sequence quality score >20, before denoising, merging of reads, removal of chimeras and clustering amplicon sequence variants (ASVs). Taxonomy was assigned using Reference database SILVA 138 release. In addition, PICRUSt v.2.0 pipeline for marker gene metagenome inference was run to predict the functional potential of the bacterial community. Further bioinformatics analyses were performed on R (v.4.4.1). Microbiome diversity indicators were calculated using microbiome (v.1.26.0) (19) and Phyloseq (v.1.48.0)(20) R packages. Ordination plots and microbiota composition bar plots were generated using microbiome (v.1.26.0) (19) R package. Statistical analysis of the microbiota profiles was carried out using several microbiome-specific statistical methods: ANCOM, ANCOMBC, ALDEx2, LEfSe and metagenomeSeq and Limma-Voom implemented in microbiomeMarker (v.1.10.0) R package (21). Sequencing reads were normalised prior to statistical analysis using total sum scaling (TSS) method implemented in microbiomeMarker (v.1.10.0) R package (21). To further investigate the effect of interindividual variability in microbiota composition profiles, linear mixed-effects models (LMMs) were computed using lme4 (v.1.1.35.5) R package. Diet and radiation were considered as fixed effects while mouse cage was considered a random effect. Post-Hoc analysis of LMM model (Tukey test) was performed using multcomp (v.1.4.26) R package. Statistically significant (*p* < 0.05 and *p_adj_* < 0.05) clades determined by both microbiomeMarker statistical methods and mixed effect model were selected as microbial biomarkers in each treatment group. Differential analysis of functional profiles predicted by PICRUSt v.2.0 was performed using ggpicrust2 v.2.5.17 R package and LinDA (linear model for differential abundance analysis of microbiome compositional data) method.

Statistically significant (*p* <0.05) correlations between microbiota composition profiles, fermentation acid levels and tumour growth rates were calculated and expressed as Pearson correlation coefficients using base R (v.4.4.1.) functions. In addition, microbiota composition and fermentation acid data were integrated using DIABLO (Data Integration Analysis for Biomarker Discovery using Latent Variable Approaches for Omics Studies) algorithm implemented in mix Omics (v.6.28.0) R package (22).

## Results

### Inulin slowed tumour growth in three prostate cancer flank models

We wanted to identify a suitable mouse prostate cancer flank allograft model, with appropriate tumour growth rates, which could be used in subsequent ionising radiation experiments with dietary fibre supplementation.

The prostate tumour models (DVL3, Myc-CaP, RM1) exhibited distinct growth kinetics (**Figure 1A–C**). DVL3 tumours grew most slowly, with 0.2% cellulose-fed mice requiring 48–55 days to reach a volume of at least 00 mm³, whereas RM-1 tumours were the most rapidly growing, reaching the same volume within 10 days.

**Figure 1.**
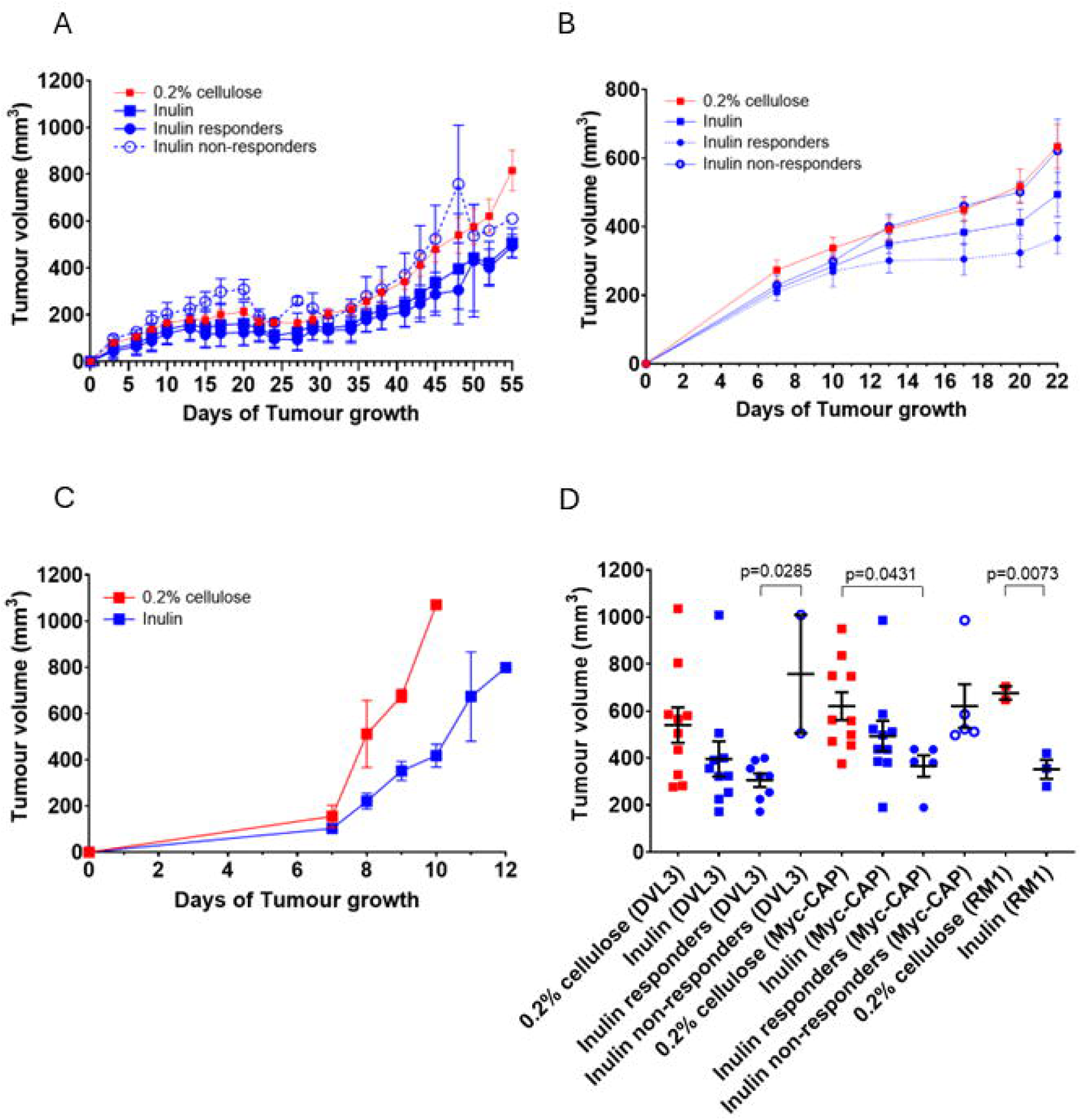
Tumour growth in mice fed 0.2% cellulose or 10% inulin diets, after injecting (A) 20 male C57BL/6J mice with 1 million prostate cancer cells (DVL3) (0.2% cellulose, n=10; 10% inulin n=10), (B) 20 male FVB mice with 1 million prostate cancer cells (Myc-CaP) (0.2% cellulose, n=10; 10% inulin n=10), (C) six male C57BL/6J mice with 0.5 million prostate cancer (RM-1) cells (0.2% cellulose, n=3; 10% inulin n=3); (D) Comparison of tumour growth at days 48, 22 and 9 for DVL3, Myc-CaP and RM-1 cells, respectively, by one-way ANOVA followed by Tukey’s multiple comparison test, excepting tumour growth of the RM-1 cells which was compared with an unpaired t-test with Welch’s correction. Data are presented as mean ± SEM.

Across all three tumour models, the average tumour growth in inulin fed mice was delayed compared to that in mice fed 0.2% cellulose, although only statistically significant for the RM-1 model (DVL3, p=0.19; Myc-CaP, p=0.14; RM-1, p=0.0073, unpaired t-test with Welch correction; **Figure 1A to C**; see **Figure S1** for individual mouse growth curves for DVL3 and Myc-CaP). However, using a tumour volume threshold at 450 mm^3^ revealed heterogeneity of responses, with some inulin-fed mice showing delayed tumour growth (responders) while others did not (non-responders; **Figure 1A, B, Figure S1**). Tumour growth comparisons were made at days 48, 22 and 9 for DVL3, Myc-CaP and RM1 cells, respectively (Figure 1D). Inulin responders showed significantly delayed tumour growth relative to non-responders in the DVL3 C57BL/6J model (p=0.029; one-way ANOVA with Tukey’s multiple comparison test), but not in the Myc-CaP FVB model (p=0.10), although in Myc-CaP mice growth was delayed in inulin responders relative to 0.2% cellulose (p=0.043).

Due to the slow rates of tumour growth, the DVL3 model was not advanced to the irradiation experiments. We also tested a Myc-CaP model in C57BL/6 mice (cells were a generous gift from Dr Leigh c, Cedars-Sinai Cancer, LA (23), as these mice are a more common, less expensive mouse model than male FVB mice. Furthermore, FVB mice tend to be more aggressive(24), often requiring retention in original cage cohorts. Unfortunately, this model was unsuccessful in our hands, so we proceeded with the Myc-CaP FVB model in our radiation experiments.

### Inulin slowed tumour growth post-IR in the Myc-CaP FVB model and impacted on the gut microbiota

We chose to investigate inulin first, for its potential applicability to future human cancer trials, as it is commonly used in human dietary studies and is well tolerated and freely available from health food shops, for its various health benefits.

In the FVB-Myc-CaP model, irradiated mice fed an inulin-enriched diet exhibited significantly delayed tumour growth at Day 12 compared with mice fed 0.2% cellulose (p=0.014 by two-way ANOVA with Tukey’s multiple comparison test; **Figure 2A**); individual tumour growth curves post-IR shown in **Figure 2B**. Following irradiation, the time required for tumours to quadruple in volume was prolonged in inulin-fed mice relative to cellulose-fed controls, although not reaching significance (p = 0.057, Log-rank test for trend; **Figure 2C**) and individual comparisons were non-significant by one-way ANOVA with Tukey’s multiple comparison test (**Figure 2D**).

**Figure 2.**
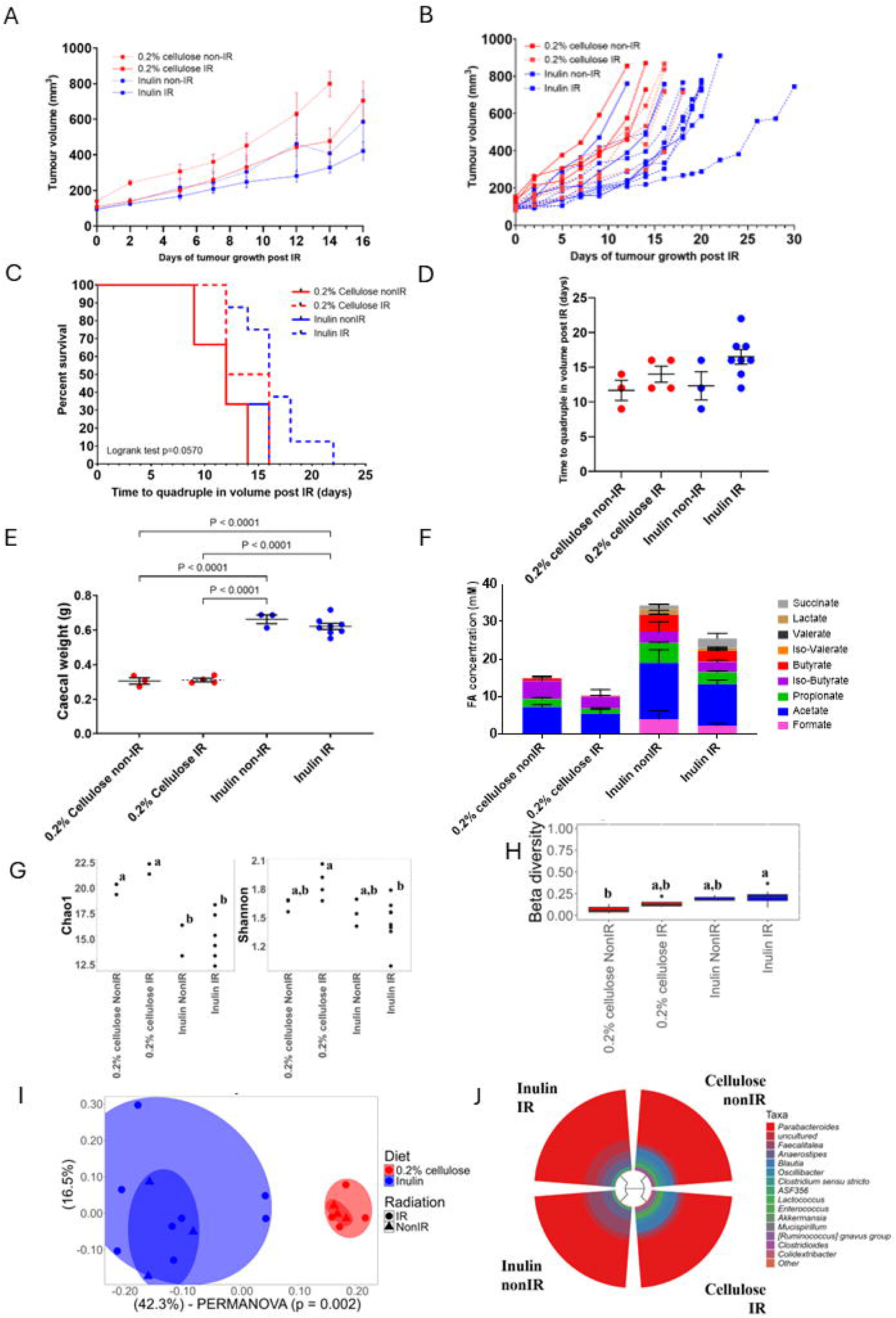
Myc-CaP tumour growth and gut microbiota analyses in male FVB mice following IR. Mice were injected with 1 million Myc-CaP prostate cancer cells and started on respective diets before tumours were irradiated to 6 Gy when they had reached 80-100 mm^3^. (A) Tumour growth post-IR (6 Gy) in mice fed 0.2% cellulose or inulin diets, (with interpolation as required; 0.2% cellulose no IR, n=3, IR, n=4; 10% inulin no IR, n=3, IR n=8). (B) Individual tumour growth curves for Figure 2A. (C) Time taken for tumours to quadruple in volume post-IR (or from size equivalent for no-IR mice), analysed by Log-rank test for trend. (D) Plot of time to quadruple in volume for individual mice, compared by one-way ANOVA with Tukey’s multiple comparison test. (E) mouse caecal weights, compared by one-way ANOVA with Tukey’s multiple comparisons test (0.2% cellulose no IR, n=3, IR, n=4; 10% inulin no IR, n=3, IR n=8). (F) Fermentation acids concentrations, compared by two-way ANOVA with Tukey’s multiple comparisons test. (G) Alpha diversity (Chao1 and Shannon indices) and (H) beta diversity analysis (Bray-Curtis dissimilarity metric) of the microbiota profiles of mice fed 0.2% cellulose or inulin diets and receiving 6 Gy irradiation (IR) or no irradiation (nonIR), ^a,b^ statistically significant (*p* <0.05) differences between groups, (I) Principal coordinates analysis (PCoA) and, (J) cladogram of the microbiota composition profiles.

The caecal weights were significantly higher in inulin-fed mice when compared to 0.2% cellulose with or without IR, and for 0.2% cellulose non-IR vs Inulin IR (p<0.001, **Figure 2E**), reflecting the fermentation of inulin within that organ in mice (25). While formate, lactate, iso-valerate, valerate and succinate were present only in the inulin-fed group, the acetate concentrations were higher in caeca from the inulin IR vs 0.2% cellulose IR (p <0.0001) and inulin non-IR vs 0.2% cellulose non-IR mice (p <0.0001), and the inulin non-IR vs inulin IR (p=0.012) groups, but this was not seen for butyrate or propionate.

Furthermore, there were no differences in iso-butyrate concentrations between these groups (**Figure 2F**). The rate of tumour growth post-IR showed no association with the concentrations of acetate or butyrate in the inulin-fed group by linear regression (**Figure S2A** and **B**).

In the Myc-CaP tumour-bearing FVB mice, microbial richness was higher by Chao1 index (p <0.05) in 0.2% cellulose-fed compared to inulin-fed mice, with no statistically significant differences between non-IR and IR groups (**Figure 2G**), and, although the Shannon index alpha diversity measure was not significantly different between 0.2% cellulose and inulin non-IR groups, the inulin IR group showed the lowest Shannon index values and was statistically significantly lower than the 0.2% cellulose IR group (p<0.05; **Figure 2G**). Inulin-fed mice showed a trend for higher, but non-statistically significant beta diversity (p=0.17) compared to 0.2% cellulose-fed mice, with no significant (*p*>0.05) differences between non-IR and IR groups (**Figure 2H**).

Principal Coordinates Analysis (PCoA) accurately discriminated samples corresponding to 0.2% cellulose and inulin diets, highlighting characteristic microbiota patterns (**Figure 2I**). The most abundant genera in the microbiota of both inulin-and 0.2% cellulose-fed FVB mice were *Parabacteroides*, *Faecalitalea*, *Anaerostipes*, *Blautia* and an uncultured genus (**Figure 2J**).

Microbiota analysis using mixed effect models identified several microbial biomarkers (**Table S2**): 0.2% cellulose-fed FVB mice showed high abundances of *Blautia* while inulin-fed mice were characterised by high abundances of an uncultured genus, *Faecalitalea and Anaerostipes* (**Table S2A**). The effect of radiation was also investigated. *Parabacteroides* abundances were high in the 0.2% cellulose non-IR group probably due to a reduction in the uncultured taxa, while *Faecalitalea* and *Anaerostipes* were characteristic of the inulin non-IR group, and radiation led to an increase in an uncultured genus in inulin group (**Table S2B**). Correlation analysis revealed positive correlations between several SCFA-producing genera (*Anaerostipes*, *Eubacterium coprostanoligenes* group and *Faecalitalea*) and the majority of fermentation acids in Myc-CaP tumour-bearing FVB mice (**Figure S3A**). Tumour growth rate was not associated with *Ruminococus gnavu* abundance in inulin-fed or 0.2% cellulose-fed mice by linear regression **(Figure S2C).** Furthermore, inulin NonIR diets led to high abundances of *Anaerostipes* and formate and propionate levels according to data integration models (**Figure S3B).**

### Inulin, pectin and **β**-glucan slowed tumour growth post-IR in the RM-1 C57BL/6J model and impacted on the gut microbiota

Having studied inulin extensively in prostate and previously in bladder cancer mouse models, we then wished to examine the effects of other fibres on the gut microbiota, which might be exploited clinically in the future.

In the fast growing C57BL6 RM-1 tumour model, there was significantly reduced tumour growth observed in β-glucan- and inulin-fed nonIR- and IR-mice compared to 0.2% cellulose non-IR mice at Day 2 post-IR (all p<0.05) by mixed effects analysis with multiple comparisons (**Figure 3A**). When comparing tumour size at three days post-IR (and equivalent for nonIR mice), by one-way ANOVA, there was a significant difference among diet groups (p=0.0015) and, by Tukey’s multiple comparisons test, differences were observed between nonIR 0.2% cellulose and nonIR β-glucan (p=0.013) and IR β-glucan (p=0.048) and between nonIR 0.2% cellulose and nonIR inulin (p=0.034) and IR inulin (p=0.0062), and between nonIR 0.2% cellulose and IR pectin (p=0.014) but not nonIR pectin (p=0.22). Tumour growth rate was also slower in inulin IR compared to 0.2% cellulose IR (p=0.017**; Figure S4**). High fibre diets delayed the time taken for tumours to quadruple in volume post-IR, by log-rank test for trend (p=0.009; **Figure 3B**), compared to 0.2% cellulose nonIR: pectin IR, p=0.016; β-glucan, p=0.013; inulin, p=0.0058. Furthermore, inulin IR was delayed compared to 0.2% cellulose IR, p=0.043 **(Figure 3C**; one-way ANOVA with Tukey’s multiple comparisons test). High fibre diets also resulted in significantly heavier caeca compared to the low fibre diet (0.2% cellulose)-fed mice (all p<0.02;**Figure 3D**).

**Figure 3.**
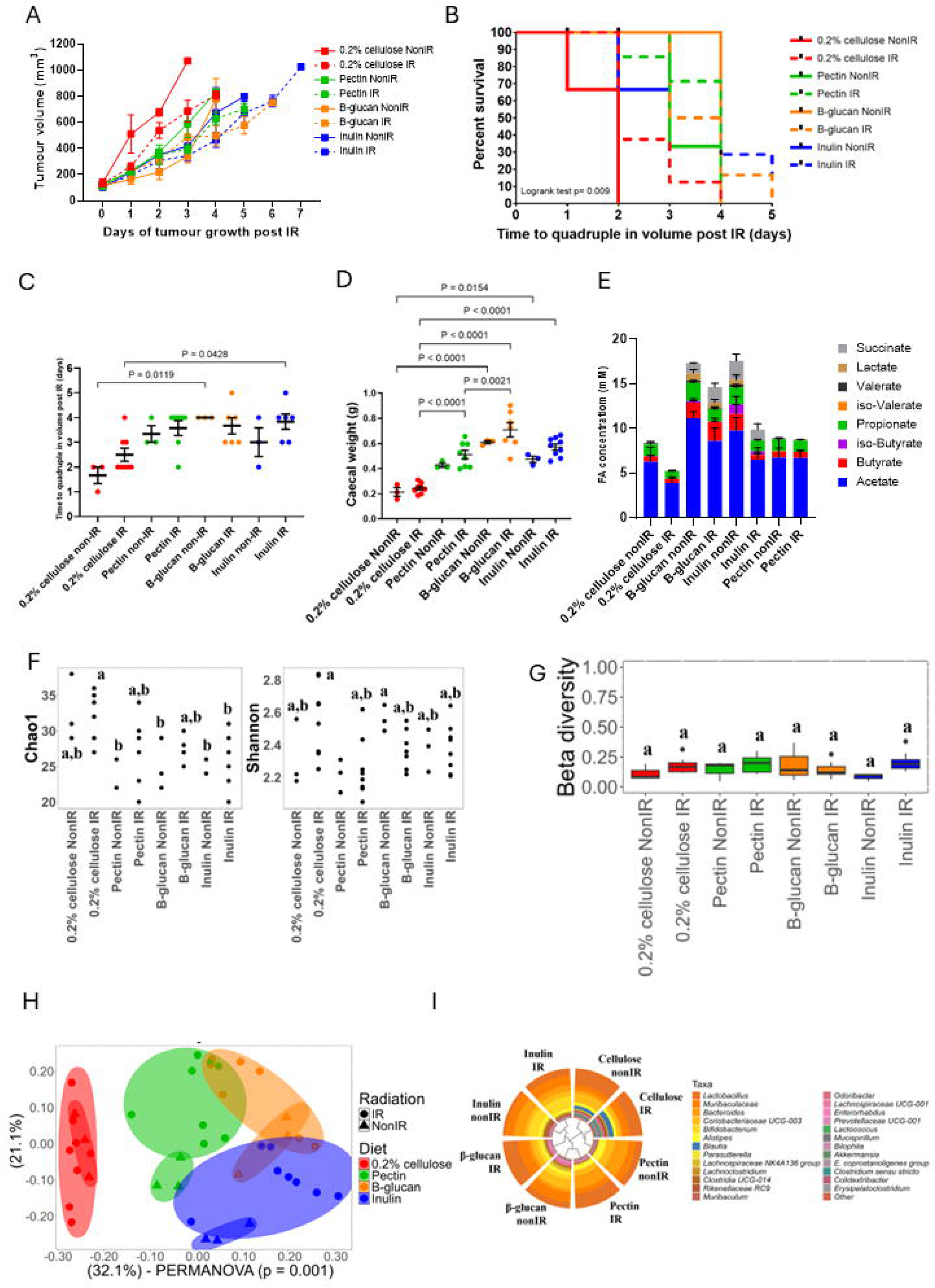
Tumour growth and microbiota analyses in male C57BL/6J mice injected with 0.5 million prostate cancer (RM-1) cells. (A) tumour growth post-IR (6 Gy) in mice fed 0.2% cellulose, pectin, β-glucan or inulin diets (0.2% cellulose IR, n=8; pectin IR, n=8; β-glucan IR, n=7; inulin IR, n=9; no IR, all n=3) compared by mixed effects analysis with multiple comparisons. (B) time taken for tumours to quadruple in volume post-IR (or from size equivalent for non-IR mice), assessed by log-rank test for trend (0.2% cellulose IR, n=8; pectin IR, n=7; β-glucan IR, n=6; inulin IR, n=6; no IR, all n=3). Five mice were censored (pectin IR, Day 1; β-glucan IR, Day 2; inulin IR, Days 1, Day 2 and Day 4) as they were culled before their tumours could quadruple in volume. (C) Plot of time to quadruple in volume for individual mice compared by one-way ANOVA with Tukey’s multiple comparisons test (0.2% cellulose no IR, n=3, IR, n=8; pectin no IR, n=3, IR, n=7; β-glucan no IR, n=3, IR, n=6; inulin no IR, n=3, IR, n=6). (D) Mouse caecal weights compared by one-way ANOVA with Tukey’s multiple comparison test. (E) Fermentation acid concentrations compared by two-way ANOVA with Tukey’s multiple comparison test. Data are presented as mean ± SEM. (F) Alpha diversity and (G) beta diversity analysis of the microbiota profiles of tumour-bearing C57BL/6J mice fed 0.2% cellulose, inulin, β-glucan or pectin diets and receiving 6 Gy irradiation (IR) or no irradiation (nonIR). (H) PCoA of the microbiota profiles of tumour-bearing C57BL/6J mice fed 0.2% cellulose, inulin, β-glucan or pectin diets and receiving 6 Gy irradiation (IR) or no irradiation (nonIR). ^a,b^ Statistically significant (*p* < 0.05) differences between groups. (I) Cladogram showing the microbiota composition profiles of tumour-bearing C57BL/6J mice fed 0.2% cellulose, inulin, β-glucan or pectin diets and receiving 6 Gy irradiation (IR) or no irradiation (nonIR).

In terms of fermentation acids, 0.2% cellulose-fed mice had no detectable lactate or succinate levels. Compared to the 0.2% cellulose IR group, acetate concentrations were significantly higher in the β-glucan, inulin and pectin IR groups (all p<0.0001) and compared to the 0.2% nonIR group, acetate concentrations were significantly higher in the β-glucan (p<0.0001) and inulin (p=0.0032) but not pectin (p=NS) nonIR groups (**Figure 3E**) but butyrate and propionate comparisons were all non-significant. Neither acetate nor butyrate concentrations were significantly associated with growth rate post-IR in inulin-fed or β-glucan-fed mice, by linear regression **(Figure S5B**(i) to (iv)).

The 0.2% cellulose IR group showed the highest Chao1 index, while no significant differences were observed in Shannon index between any of the groups (**Figure 3F**). There were no significant differences in beta-diversity (*p*=0.20) between diets (0.2% cellulose, inulin, pectin and β-glucan) or between non-IR and IR groups (*p*>0.05; **Figure 3G**). Using Principal Coordinates Analysis (PCoA). different groups of samples were observed for each diet (**Figure 3H**). In general, samples were clustered by diet regardless of irradiation status, highlighting the resilience of fibre-degrading microbiota in both the C57BL/6J model (**Figure 3H**) and in the FVB model (**Figure 2I**), and also reflecting the relatively localised 1.5 cm applicator radiotherapy field.

In the C57BL6 RM-1 model, the predominant microbial genera were *Lactobacillus,* an unknown *Muribaculaceae* genus, *Bacteroides*, novel *Coriobacteriaceae* genus UCG-003, *Bifidobacterium and Alstipes*. Mice fed 0.2% cellulose additionally showed high *Blautia* abundances while inulin promoted *Bifidobacterium* (**Figure 3I**; **Table S3A**). *Bacteroides* and *Alistipes* were enriched in pectin groups while *Lactobacillus* and an unknown *Muribaculaceae* genus were characteristic of β-glucan-fed mice (**Table S3A**). The statistically significant effects of IR were also investigated. Of note, the IR pectin group showed higher *Prevotellaceae* UCG-001 abundances compared to the non-IR pectin group. Similarly, several *Lachnospiraceae* members including *Lachnoclostridium* and *Lachnospiraceae* UCG-001 were enriched in β-glucan non-IR compared to β-glucan IR (**Table S3B**).

Of note, microbiota composition bar plots suggest that the effects of diet were greater than effects of cages on the microbiota profiles (**Figure S6**). PERMANOVA analysis revealed no significant (p > 0.05) differences between cages corresponding to pectin, β-glucan and inulin diet groups.

Correlation analysis revealed positive correlations of *Anaerostipes*, *Bifidobacterium*, *Lachnospiraceae* NK4A13 and *Muribaculaceae* abundances with several fermentation acids (**Figure S7A**). Several bacterial genera (*Akkermansia*, *Bifidobacterium*, *Eubacterium coprostanoligene*, *Lachnospiraceae* NK4A13 and *Muribaculum*) showed negative correlations with tumour growth rates in the C57BL6 RM-1 model (**Figure S7B**). Notably, β-glucan diets (non-IR and IR groups) were characterised by an enrichment in *Akkermansia*, *Lachnospira* members and high butyrate and lactate and total fermentation acid concentrations (**Figure S7C**).

### Responders vs non-responders to dietary fibre and IR

We observed throughout our models, that tumours in mice fed inulin grew at varying rates and could be dichotomised into responders vs non-responders (C57BL/6J mice bearing DVL3 tumours in **Figures 1A** and **S1A**; FVB mice bearing Myc-CaP tumours in **Figures 1B** and **S1B**), with reduced tumour growth in responders compared to non-responders. Moreover, when using a cut off 400 mm^3^ at Day 16 post-IR for inulin-fed, irradiated C57BL/6J mice bearing DVL3 tumours (**Figure 4A**) data were dichotomised into five responders and three non-responders (p=0.020; one-way ANOVA, with Tukey’s multiple comparisons test). In the RM-1 irradiated mice (**Figure 4B (i)-(iv)**), pectin, β-glucan and inulin-fed, but not 0.2% cellulose-fed, mice showed a dichotomous effect at Day 3 post-IR with the 400 mm^3^ cut off, but this was only significant for β-glucan (p=0.029) and inulin-fed mice (p=0.0082), by one-way ANOVA, with Tukey’s multiple comparisons test. There were no obvious cage effects seen in terms of responder vs non-responder status (**Table S4**), and no significant differences by two-way ANOVA.

**Figure 4.**
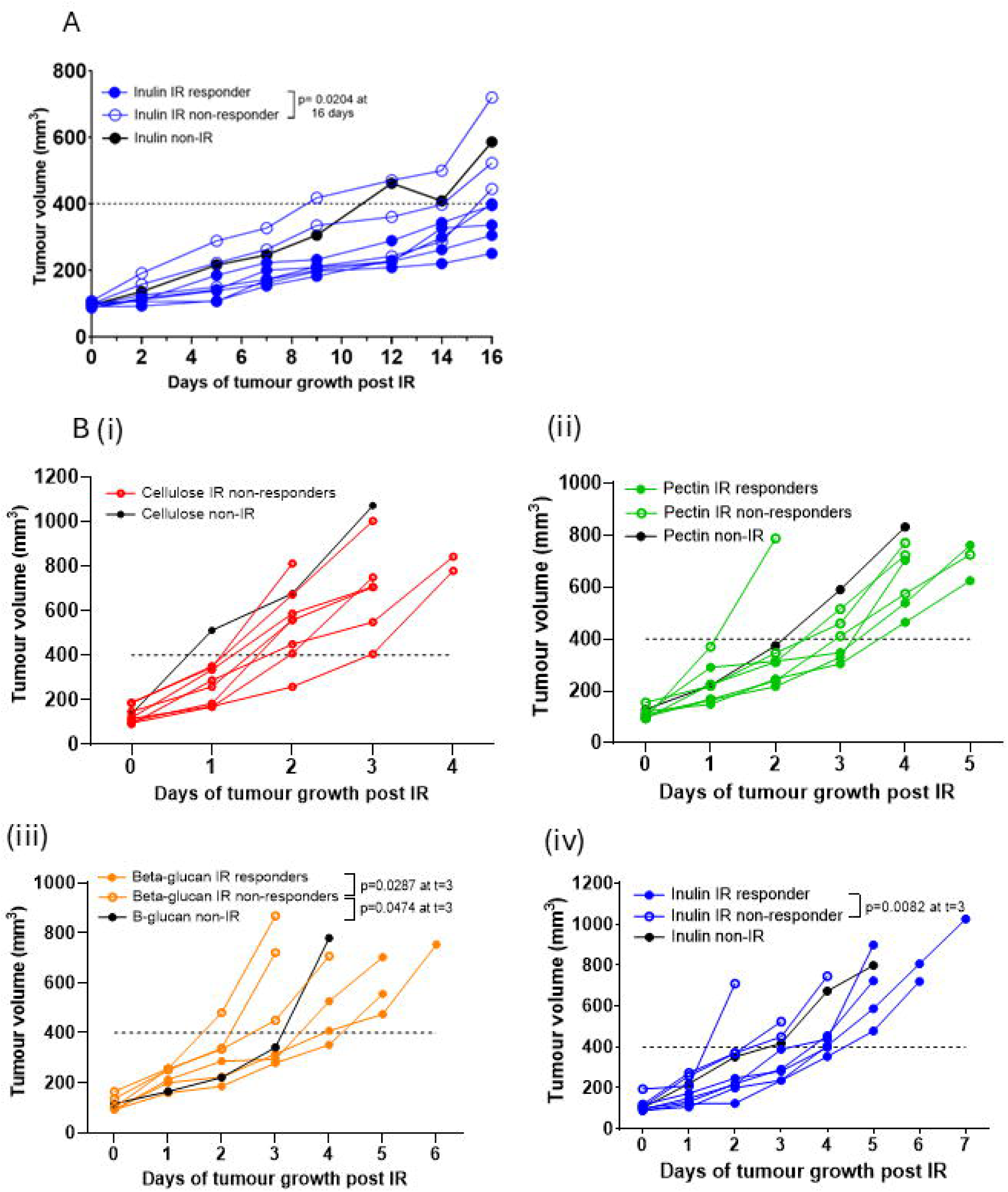
Responders and non-responders to IR and dietary interventions in Myc-CaP FVB mice and RM-1 C57BL/6J mice. (A) Comparison of responders and non-responders to IR and inulin in Myc-CaP FVB mice (responder, n=5; non-responder, n=3) by one-way ANOVA with Tukey’s multiple comparison test at Day 16). Responders were classified as a tumour growth of <400 mm^3^ by Day 16 post IR. Non-IR plot of mean of n=3 is shown for visual reference. Inulin IR responders vs non-responders showed a significant difference (p=0.0204). (B) Comparison of RM-1 C57BL/6J mice responders and non-responders to IR and (i) 0.2% cellulose (non-responder, n=8), (ii) pectin (responder, n=3, non-responder, n=4), (iii) β-glucan (responder, n= 3, non-responder, n=3) and (iv) inulin (n=5, non-responder, n=3). Responders were classified as a tumour growth <400 mm^3^ by Day 3. In all non-IR groups, n=3 and mean data are presented.

Differences in microbiota composition between responders and non-responders among the inulin-, pectin- and β-glucan-fed mice were determined (relative abundances **Table 1**). Responder inulin-fed Myc-CaP FVB mice were characterised by high relative abundances of *Parabacteroides* and *Faecalibaculum*, while responder inulin-fed RM-1 C57BL/6J mice were characterised by high abundances of *Bifidobacterium*. On the other hand, responder pectin-fed RM-1 C57BL/6J mice showed high abundances of *Lactobacillus* and *Parasutterella,* while responder β-glucan-fed mice were characterised by high abundances of *Muribaculum* and an unknown *Muribaculaceae* genus (**Table 1**). It should be noted that *Muribaculaceae* and *Lachnospiraceae* are predominant bacterial families in healthy mouse gut microbiota (26).

**Table 1.**
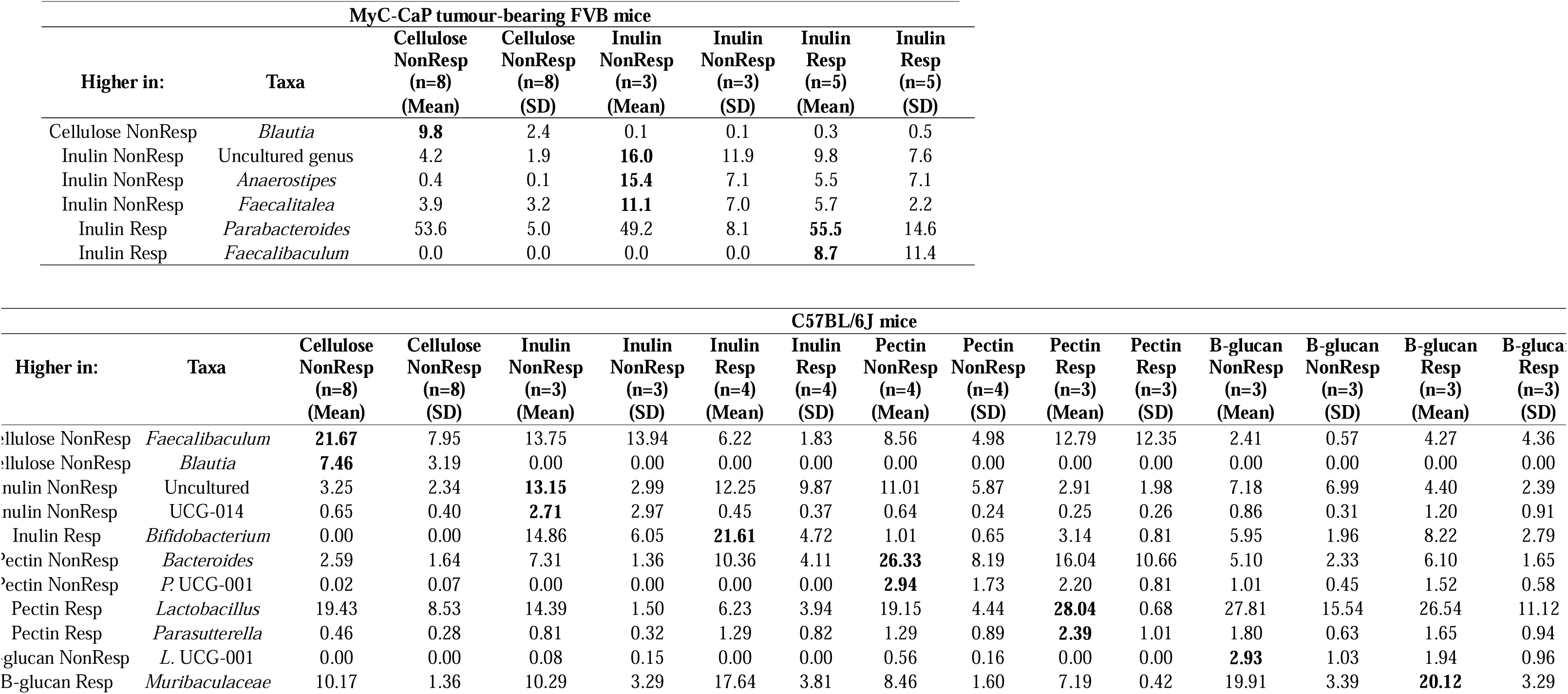
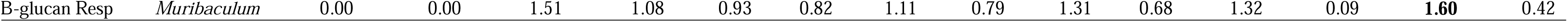
Microbial genera showing high abundance in the taxonomic profiles and identified as microbial biomarkers of responder (Resp) and non-responder (NonResp) MyC-CaP tumour-bearing FVB mice fed 0.2% cellulose or inulin diets (top), and RM-1 tumour bearing C57BL/6J mice fed 0.2% cellulose, inulin, β-glucan or pectin diets (bottom). Data are expressed as sequence abundance percentage (%). SD: standard deviation. ***L.* UCG-001**: *Lachnospiraceae* UCG-001, ***P.* UCG-001**: *Prevotellaceae* UCG-001, **UCG-014**: *Clostridia* UCG-014. Potential microbial biomarkers (*p* <0.05 and *p_adj_* <0.05) of each group determined by microbiomeMarker statistical methods are highlighted in bold.

| MyC-CaP tumour-bearing FVB mice |  |  |  |  |  |  |  |
| --- | --- | --- | --- | --- | --- | --- | --- |
| Higher in: | Taxa | Cellulose<br>NonResp<br>(n=8)<br>(Mean) | Cellulose<br>NonResp<br>(n=8)<br>(SD) | Inulin<br>NonResp<br>(n=3)<br>(Mean) | Inulin<br>NonResp<br>(n=3)<br>(SD) | Inulin<br>Resp<br>(n=5)<br>(Mean) | Inulin<br>Resp<br>(n=5)<br>(SD) |
| Cellulose NonResp | *Blautia* | **9.8** | 2.4 | 0.1 | 0.1 | 0.3 | 0.5 |
| Inulin NonResp | Uncultured genus | 4.2 | 1.9 | **16.0** | 11.9 | 9.8 | 7.6 |
| Inulin NonResp | *Anaerostipes* | 0.4 | 0.1 | **15.4** | 7.1 | 5.5 | 7.1 |
| Inulin NonResp | *Faecalitalea* | 3.9 | 3.2 | **11.1** | 7.0 | 5.7 | 2.2 |
| Inulin Resp | *Parabacteroides* | 53.6 | 5.0 | 49.2 | 8.1 | **55.5** | 14.6 |
| Inulin Resp | *Faecalibaculum* | 0.0 | 0.0 | 0.0 | 0.0 | **8.7** | 11.4 |

| C57BL/6J mice |  |  |  |  |  |  |  |  |  |  |  |  |  |  |  |
| --- | --- | --- | --- | --- | --- | --- | --- | --- | --- | --- | --- | --- | --- | --- | --- |
| Higher in: | Taxa | Cellulose<br>NonResp<br>(n=8)<br>(Mean) | Cellulose<br>NonResp<br>(n=8)<br>(SD) | Inulin<br>NonResp<br>(n=3)<br>(Mean) | Inulin<br>NonResp<br>(n=3)<br>(SD) | Inulin<br>Resp<br>(n=4)<br>(Mean) | Inulin<br>Resp<br>(n=4)<br>(SD) | Pectin<br>NonResp<br>(n=4)<br>(Mean) | Pectin<br>NonResp<br>(n=4)<br>(SD) | Pectin<br>Resp<br>(n=3)<br>(Mean) | Pectin<br>Resp<br>(n=3)<br>(SD) | B-glucan<br>NonResp<br>(n=3)<br>(Mean) | B-glucan<br>NonResp<br>(n=3)<br>(SD) | B-glucan<br>Resp<br>(n=3)<br>(Mean) | B-glucan<br>Resp<br>(n=3)<br>(SD) |
| Cellulose NonResp | *Faecalibaculum* | **21.67** | 7.95 | 13.75 | 13.94 | 6.22 | 1.83 | 8.56 | 4.98 | 12.79 | 12.35 | 2.41 | 0.57 | 4.27 | 4.36 |
| Cellulose NonResp | *Blautia* | **7.46** | 3.19 | 0.00 | 0.00 | 0.00 | 0.00 | 0.00 | 0.00 | 0.00 | 0.00 | 0.00 | 0.00 | 0.00 | 0.00 |
| Inulin NonResp | Uncultured | 3.25 | 2.34 | **13.15** | 2.99 | 12.25 | 9.87 | 11.01 | 5.87 | 2.91 | 1.98 | 7.18 | 6.99 | 4.40 | 2.39 |
| Inulin NonResp | UCG-014 | 0.65 | 0.40 | **2.71** | 2.97 | 0.45 | 0.37 | 0.64 | 0.24 | 0.25 | 0.26 | 0.86 | 0.31 | 1.20 | 0.91 |
| Inulin Resp | *Bifidobacterium* | 0.00 | 0.00 | 14.86 | 6.05 | **21.61** | 4.72 | 1.01 | 0.65 | 3.14 | 0.81 | 5.95 | 1.96 | 8.22 | 2.79 |
| Pectin NonResp | *Bacteroides* | 2.59 | 1.64 | 7.31 | 1.36 | 10.36 | 4.11 | **26.33** | 8.19 | 16.04 | 10.66 | 5.10 | 2.33 | 6.10 | 1.65 |
| Pectin NonResp | *P. UCG-001* | 0.02 | 0.07 | 0.00 | 0.00 | 0.00 | 0.00 | **2.94** | 1.73 | 2.20 | 0.81 | 1.01 | 0.45 | 1.52 | 0.58 |
| Pectin Resp | *Lactobacillus* | 19.43 | 8.53 | 14.39 | 1.50 | 6.23 | 3.94 | 19.15 | 4.44 | **28.04** | 0.68 | 27.81 | 15.54 | 26.54 | 11.12 |
| Pectin Resp | *Parasutterella* | 0.46 | 0.28 | 0.81 | 0.32 | 1.29 | 0.82 | 1.29 | 0.89 | **2.39** | 1.01 | 1.80 | 0.63 | 1.65 | 0.94 |
| β-glucan NonResp | *L. UCG-001* | 0.00 | 0.00 | 0.08 | 0.15 | 0.00 | 0.00 | 0.56 | 0.16 | 0.00 | 0.00 | **2.93** | 1.03 | 1.94 | 0.96 |
| B-glucan Resp | *Muribaculaceae* | 10.17 | 1.36 | 10.29 | 3.29 | 17.64 | 3.81 | 8.46 | 1.60 | 7.19 | 0.42 | 19.91 | 3.39 | **20.12** | 3.29 |
| B-glucan Resp | *Muribaculum* | 0.00 | 0.00 | 1.51 | 1.08 | 0.93 | 0.82 | 1.11 | 0.79 | 1.31 | 0.68 | 1.32 | 0.09 | **1.60** | 0.42 |

To gain a better understanding of the potential activities of microbiome profiles of responders and non-responders, functional predictions were generated using PICRUSt v.2.0 software (**Figures S8-11**). Differences in functional profiles were diet driven, with no major differences between non-responders and responders to diet and irradiation within specific diet groups. In the Myc-CaP FVB mice, predicted functional activities of the microbiota profiles revealed higher abundances of lipoarabinomannan, toluene, ethylbenzene and steroid metabolism in the non-responder 0.2% cellulose diet group compared to the non-responder inulin group (**Figure S8**).

In terms of RM-1 C57BL/6J mice pectin and β-glucan responders, the β-glucan diet group showed higher abundances of metabolic pathways involved in cell function and interactions compared to the pectin group (**Figure S9**). When comparing inulin versus pectin non-responder diet groups, lipoarabinomannan and unsaturated fatty acid metabolic pathways were seen to be more abundant in the inulin diet group compared to the pectin group (**Figure S10**). Glycan biosynthesis, lipid metabolism and xenobiotics biodegradation pathways showed higher abundances in the non-responder β-glucan diet group compared to the non-responder pectin group (**Figure S11**).

### Normal tissue response to ionising radiation

To determine the effects of the various fibres on intestinal normal tissue responses to IR, the intestinal crypt assay was used (15,16), Initial morphological analysis in C57BL/6J mice showed that increasing radiation dose from 10 to 14 Gy increased the damage to intestinal integrity, characterised by the blunting of the villi and reduced numbers of regenerating crypts across all fibre types. (**Figure 5A**). Further quantitative analysis found that, as the dose of ionising radiation increased, the percentage of surviving crypts decreased in all dietary groups, with a significant difference between 0.2% inulin and inulin at 12 Gy (p=0.043), by two-way ANOVA with Tukey’s multiple comparisons test (**Figure 5B**). At 12 Gy, the percentage of surviving crypts was significantly higher in mice fed inulin (p=0.032), and although a trend was observed for pectin (p=0.074) and β-glucan (p=0.064) this did not reach statistical significance by one-way ANOVA with Tukey’s multiple comparisons test (**Figure 5C**).

**Figure 5.**
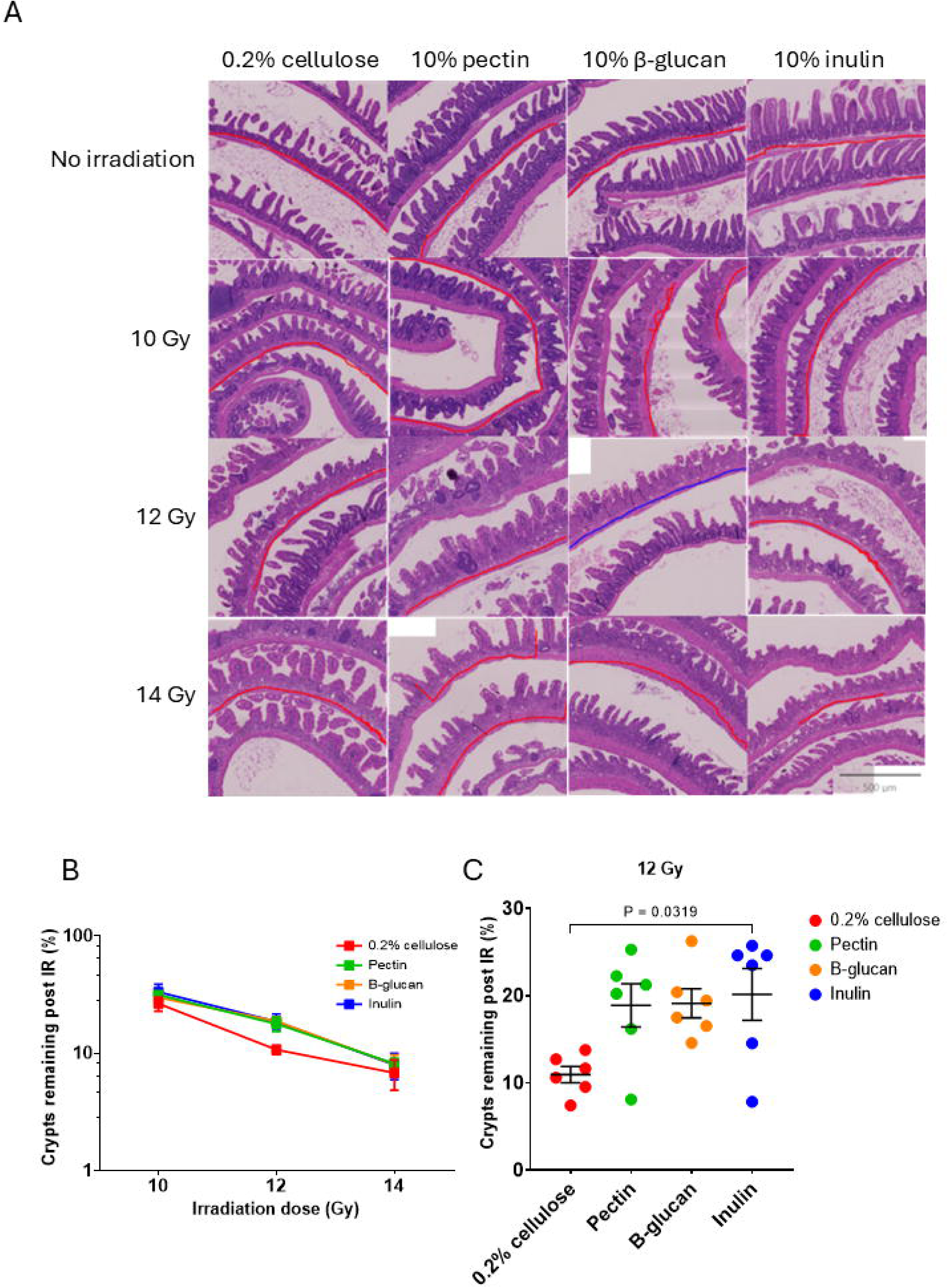
Normal tissue responses to IR and diet. (A) Representative histopathological images of gut Swiss rolls for different diets and no IR or 10, 12 or 14 Gy IR. Red line indicates a length of intestine which was used for regenerating crypt scoring. (B) Percentage of remaining crypts following irradiation in mice fed 0.2% cellulose, pectin, β-glucan and inulin, compared to respective means of n=4 no IR controls. (C) percentages of surviving crypts at 12 Gy of irradiation in mice fed 0.2% cellulose, pectin, β-glucan and inulin, compared by one-way ANOVA with Tukey’s multiple comparisons test (n=6 for all)

## Discussion

Prostate cancer is a major public health concern in men globally. Increased prostate-specific antigen screening and longer life expectancies in men have resulted in a substantial increase in the incidence and prevalence of prostate cancer in older men (4). Previous studies have reported that dietary fibre may play a role in reducing the risk of developing certain types of cancer and suggested integrating dietary strategies in comprehensive cancer care (9,27). This study examined the impact of dietary fibre in modulating the gut microbiota in murine models, and its effects on improving outcomes following ionising radiation, in terms of tumour responses and gastrointestinal toxicity.

Having previously undertaken work in mouse bladder cancer flank allografts, we aimed to identify a mouse prostate cancer allograft model, suitable for future diet and radiation studies. In addition to the models presented here, we attempted to study Myc-CaP in C57BL/6J mice (23), as using this mouse strain would have been more widely applicable than studying FVB mice which tend to be more aggressive, but tumour growth was unsuccessful in our hands.

### Antitumour effects of fibre

Across our three prostate cancer models (Myc-CaP in FVB mice, RM-1 in C57BL/6J mice, and DVL3 in C57BL/6J mice), tumour growth rate varied, with RM-1 tumours progressing most rapidly and DVL3 the slowest. Regardless of prostate cancer model and mouse strain, the high-fibre diet 10% inulin consistently showed delayed tumour growth relative to low-fibre (0.2% cellulose) controls. All three high-fibre diets (inulin, pectin and β-glucan) similarly slowed growth of RM-1 tumours and prolonged the time taken for tumours to quadruple in volume following irradiation, suggesting a common effect of fermentable fibres. However, this was a very rapidly growing model, with times to quadruple volume post-IR of only 2-5 days across models (mostly 3-4 days); a more slowly growing model may have provided more granularity in terms of comparing the three high fibre types.

We used 0.2% cellulose for our low fibre controls, rather than normal chow’, which contains variable amounts of fibre. However, in our previous studies in bladder cancer, we found that high insoluble fibre resulted in similar tumour growth rates to low fibre 0.2% cellulose (13) and 10% psyllium and normal chow resulted in tumour growth rates similar to 0.2% cellulose (11), implying that the intrinsic properties of the fermentable, complex carbohydrates inulin, pectin and β-glucan have more of an impact on tumour growth than the more insoluble fibres. Low fibre diets can result in proteolytic metabolism in the distal colon, due to lack of substrates for saccharolytic fermentation, leading to production of harmful bacterial products which are associated with increased inflammation and increased gut permeability (28). However, we did not detect significantly higher levels of the branched chain fatty acids iso-butyrate and iso-valerate in 0.2% cellulose-fed mice in this study.

The anti-tumour activity observed with the high fibre diets (inulin, pectin and β-glucan) was associated with larger caecal weights, reflecting increased bulk of intestinal matter, and higher caecal acetate concentrations in inulin and β-glucan groups. In breast and bladder cancer models, inulin and inulin plus psyllium have also shown similar effects in delaying tumour growth, which were attributed to increased SCFAs (particularly butyrate and propionate) and changes in gut microbial composition (11,29).

However, in these prostate cancer models, we saw no correlations between tumour growth rate and caecal butyrate or acetate concentrations, in contrast to our own previous findings (11) and reports that SCFAs can exert anti-cancer properties, through regulation of immune responses, radiosensitisation and epigenetic modulation of tumour cells (29–31). Our current findings could be due to the relatively rapid tumour growth in the RM-1 model, as we did see higher caecal acetate levels in the inulin and β-glucan-fed mice.

We studied fermentation acids, including SCFAs and branched chain fatty acids, but also lactate and succinate. The latter are commonly produced by gut bacteria and often used as intermediates or for cross feeding. For example, in humans, *Anaerostipes* can use lactate to form butyrate, *Phasarcolarctobacterium* spp. may metabolise succinate to form propionate, with *Bacteroides* and *Prevotella* spp, the main producers of succinate in the gut. However, excepting a significant difference between succinate levels for 0.2% cellulose and β-glucan (p=0.0499), we found no significant associations for lactate or succinate.

### Gut microbiota responses to dietary fibre and ionising radiation

Dietary fibre significantly altered the gut microbiota of tumour-bearing mice. Inulin enriched SCFA-producing genera (32), including *Anaerostipes* (butyrate)*, Bifidobacterium* (acetate) and *Faecalitalea,* whereas 0.2% cellulose favoured *Blautia* (acetate). In contrast, pectin promoted fibre-degrading genera *Bacteroides* (acetate, propionate) and *Alistipes* (acetate, propionate) while β-glucan enriched *Lactobacillus* (lactate producer)*, Lachnospiraceae* UCG-001, *Coriobacteriaceae* UCG-003 and an unknown *Muribaculaceae* genus. Irradiation had no major impact on alpha or beta diversity of different groups and preserved clustering by diet, indicating resilient fibre-associated taxa and potentially reflecting the localised irradiation area. Negative correlations between *Akkermansia, Bifidobacterium, Eubacterium* and *Lachnospiraceae* taxa and tumour growth support the role of microbial metabolites in modulating anti-tumour effects (31,33).These data suggest that fermentable fibres promote growth of bacterial communities capable of producing bioactive metabolites that suppress tumour progression and enhance response to irradiation.

Some of the differentially abundant metabolic pathways determined by PICRUSt v.2.0 could be of special interest in the context of prostate cancer. Changes to glycans are common in prostate cancer, including increased branching of complex *N-*glycans (34), fatty acid metabolism genes have been proposed as biomarkers for prostate cancer prognosis (35) and, finally, lipoarabinomannan acts as a ligand that activates Toll-like receptors TLR2 (36). In prostate cancer, this TLR activation can either promote inflammation-driven cancer progression or trigger an anti-tumour response depending on the tumour’s environment (36,37).

In our 16S rRNA gene sequencing, we chose to study the V1V2 region rather than V3V4 to provide us with higher resolution of keystone taxa relevant to our study design. The V1V2 region has superior taxonomic resolution for many *Bifidobacterium* species, unlike V3V4 which often under-represents specific Actinobacteria, including many *Bifidobacterium* spp. Furthermore, Bifidobacterium is widely recognised as the most consistently increased genus and considered a primary responder to high dietary fibre intake, including inulin and resistant starches (38).

### Responders versus non-responders

The presence of responders and non-responders among the β-glucan- and inulin-fed mice highlights inter-individual heterogeneity. One might expect with identical diets and genetic backgrounds, mice purchased from established breeders would have similar responses to diet/IR combinations. One factor which could influence the gut microbiota and hence responses would be the cage in which the mouse is living. However, we found no influence of cage effects in terms of the proportion of responders and non-responders within each cage.

### Normal tissue responses to IR and fibre

The histopathological analyses provided insights into the protective capabilities of dietary fibres on the gut when exposed to increasing doses of ionising radiation. The crypt assay showed a clear difference between the high fibre dietary groups and the low fibre group, with high fibre diets providing protection for crypts against radiation, particularly at 12 Gy of radiation. We have previously reported similar findings where mice on high fibre diets preserved crypts following radiation exposure (11), although in this previous dietary study the effects were more marked at 14 Gy rather than 12 Gy, while our previous paper on panobinostat as a radiosensitiser, effects were more marked at 12 Gy with limited differences at 10 and 14 Gy (39). This could reflect differences in mouse strains or delivery of ionising radiation. Others have more recently shown that two short-chain inulins protect against radiation-induced intestinal injury in mice irradiated with 17 Gy to the abdomen, with differential protective effects related to intestinal immunity, microbial metabolite modulation and the gut microbiota, including *Muribaculum* and *Lachnospiraceae* (40). These had higher relative abundance in β-glucan-fed compared to 0.2% cellulose-fed mice in our tumour studies. Another group created a colon-retentive inulin gel for use pre-and post-abdominal radiation in mice which increased the relative abundance of *Lachnospiraceae, Akkermansia* and *Blautia* and SCFA metabolites, with increased secretion of anti-inflammatory cytokines and, in an orthotopic colorectal cancer model, the oral inulin gels followed by 10 Gy total abdominal IR improved radiotherapy efficacy and selectively protected normal intestinal tissues by crypt assay at 3.5 days (41).

Our current study confirms the dual effects on the radiotherapy therapeutic ratio seen by Lui et al (41) and in our previous work (11), where dietary fibres improve tumour responses to radiotherapy at the same time as reducing intestinal side effects. This contrasts with radioprotectors which reduce side effects but have no impact on tumour control, and radiosensitisers which improve tumour control but often at the expense of worsening side effects.

### Limitations of the study

Our radiation doses do not directly match those seen in clinical practice where smaller multiple fractions of radiation are given. However, we chose a single 6 Gy fraction for tumour irradiation in line with our previous studies, to minimise potential harmful effects to the mice of repeated anaesthetics and irradiations, and to be a suitable dose where we can see some effects of radiation but not such a high dose that we cannot see the impact of the fibres on tumour responses. The 10, 12 and 14 Gy doses chosen for the crypt assay are in line with the published literature (42).

Our findings in mouse flank models may have limited translatability to human prostate cancer, but orthotopic models are a greater burden for the mice, and our studies give an early read out of the potential benefits of adding dietary fibre to irradiation in mouse prostate cancer cell-derived tumours. We also accept that mouse models have inherent limitations in representing the human gut microbiota.

Further work is required to understand the underlying mechanisms of the effects of the various fibres with ionising radiation. Such studies could include faecal microbiota transfer, gavage of individual bacteria and antibiotic depletion experiments to study the impact of the gut microbiota, direct SCFA supplementation and untargeted metabolomics with validation of likely targets.

## Conclusions

High-fibre diets (inulin, pectin and β-glucan) reduced tumour growth and enhanced radiotherapy response across murine prostate cancer models, associated with some higher abundances of SCFA-producing taxa and elevated concentrations of SCFAs. The high fibre diets also protected intestinal crypts from irradiation. The existence of responder versus non-responder phenotypes emphasises the importance of host-dependent variability in fibre metabolism or baseline microbiota composition in therapeutic efficacy. Our hypothesis-generating findings should be followed up by further mechanistic work investigating the role of the gut microbiota and resulting metabolites and their roles in tumour responses to dietary fibre supplementation.

Findings may have relevance to men with prostate cancer on active surveillance pathways and receiving prostate radiotherapy, with fermentable fibres having the potential to serve as low-cost, accessible treatments.

## Author responsible for Statistical analysis

Dr Carlos Sabater

## Acknowledgements

We would like thank Dr Jin Pu and Dr Zeynab Heidari (CGEBM, University of Aberdeen), for sequencing the DNA samples, Donna Henderson (Rowett Analytical, University of Aberdeen) for running the SCFA analysis, Andrea Holme (Iain Fraser Cytometry Centre, University of Aberdeen) for her expert help with running the cytokine samples on the Luminex instrument and Dr Graham Horgan for his expert statistical advice. We acknowledge histology support from NHS Grampian Biorepository, Aberdeen Royal Infirmary, Aberdeen, UK. We would like to thank Dr. Jia-Yu Ke (Research Diets, Inc.) for formulation of the mouse diets and Beneo for provision of the Orafti®β-Fit flour. Anne Kiltie is the Friends of ANCHOR Clinical Chair in Oncology, University of Aberdeen.

Rights retention statement: For the purpose of open access, the author has applied a CC BY Creative Commons Attribution (CC BY) licence to any Author Accepted Manuscript arising from this submission [we have been asked to add this by our Institution for the purposes of REF29].

## Authors’ Contributions

AM and SMH performed the animal experiments, collected and processed mouse samples. AM, AP and CS performed the analysis and interpretation of the data and drafted the manuscript. DMcC provided advice and supervision on H&E analysis of intestinal Swiss rolls. ESRCD supervised the faecal bacterial sequencing and AM, CS and MAvdH analysed the data. AM, HMW and AEK obtained grant funding. AEK and HMW supervised the work. AEK reviewed and revised the first draft of the manuscript. All authors reviewed and revised the manuscript and approved the final version.

## Data availability statement for this work

Raw sequence reads have been deposited in the Short Reads Archive (SRA) of the National Center for Biotechnology Information (NCBI) under BioProject accession number PRJNA1390088.

## Declaration of generative AI and AI-assisted technologies in the manuscript preparation process

Generative AI was not used in this manuscript.

