## Supplementary Material for "Influence of high fibre diets on the gut microbiota, prostate tumour growth and normal tissue toxicity following ionising radiation"

### High fibre diets and radiotherapy outcomes

#### Supplementary Material

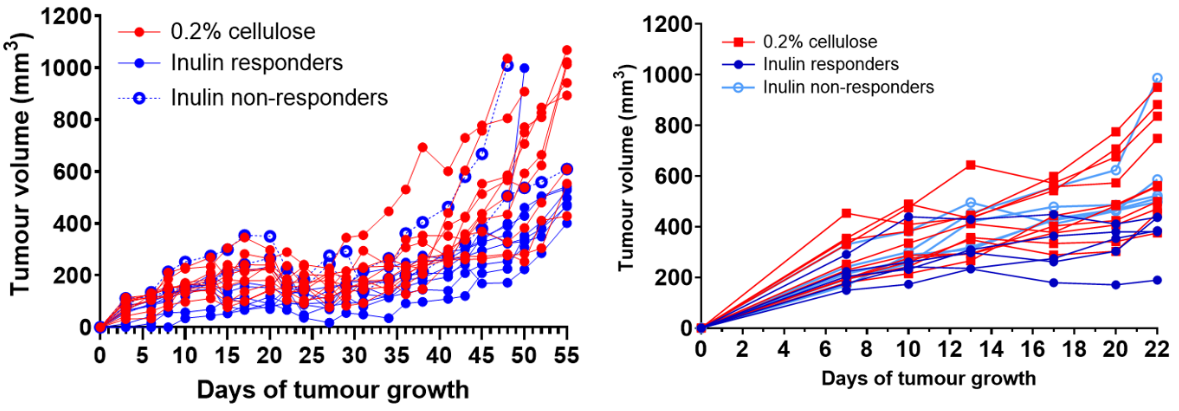

**Figure S1: Individual mouse growth curves,** in (A) DVL3 tumour model from Fig 1A (0.2% cellulose no IR, n=10, 10% inulin no IR, n=10). (B) Myc-CaP tumour model from Fig 1B (0.2% cellulose no IR, n=10, 10% inulin no IR, n=10).

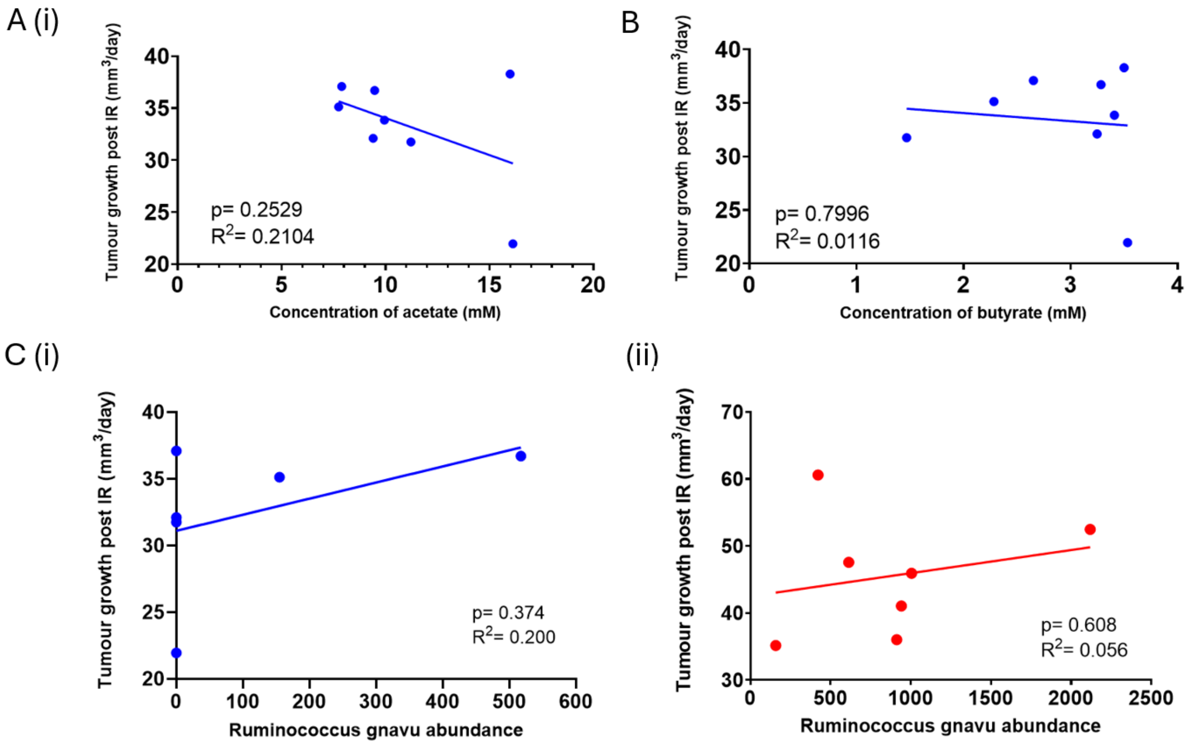

**Figure S2: Tumour growth rate post-IR in Myc-CaP FVB mice** did not correlate with (A) acetate or (B) butyrate levels in inulin-fed mice but correlated with (C) *Ruminococcus gnavus* relative abundance for (i) inulin and (ii) 0.2% cellulose-fed mice**.** Correlations by linear regression.

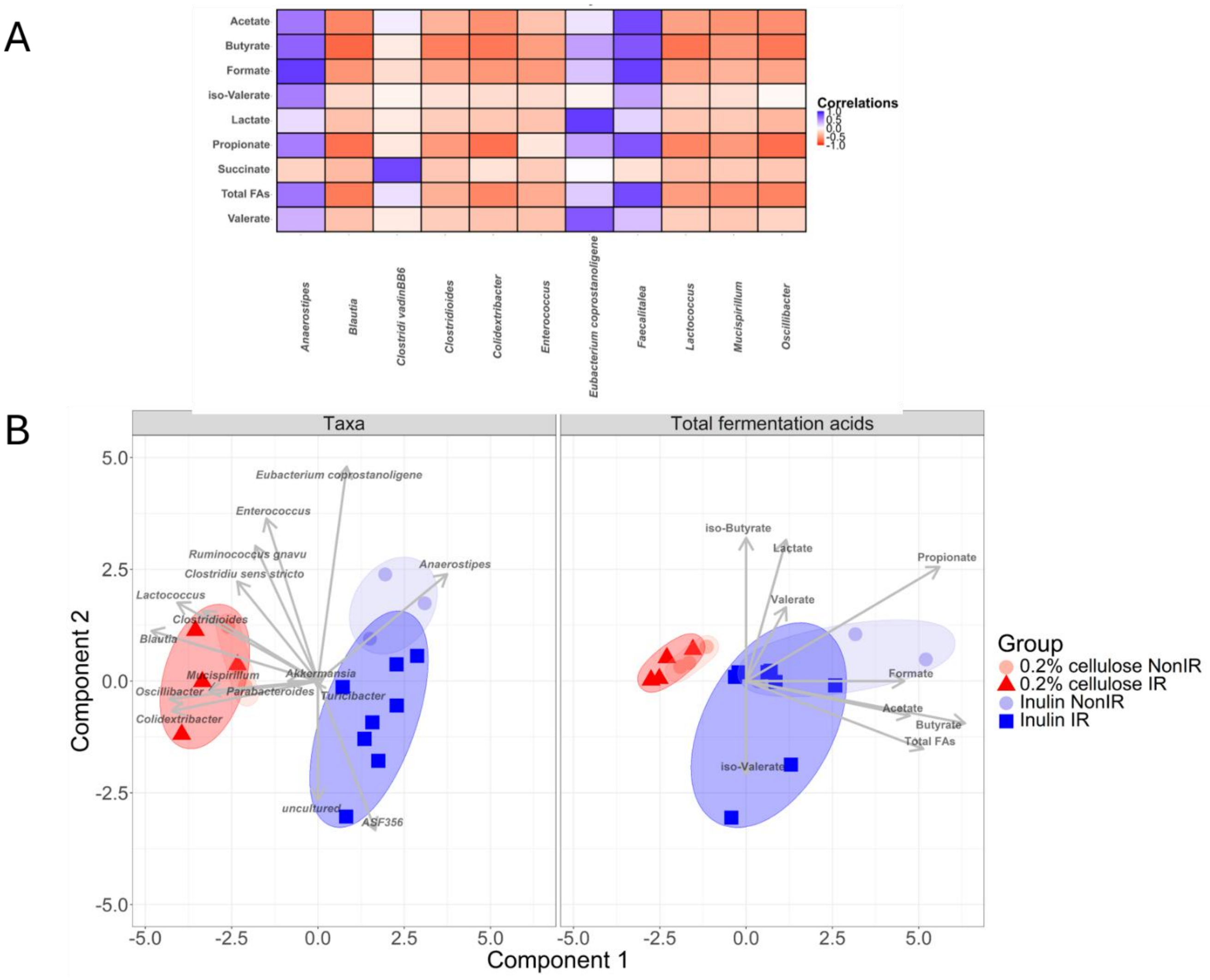

**Figure S3: Correlation heatmaps for Myc-CaP FVB mice,** showing the associations between relative abundance of specific taxa identified as the dominant/ signature taxon in one of the dietary groups (See panel C and Table S2**)** and (A) total fermentation acid levels in MyC-CaP tumour-bearing FVB mice fed 0.2% cellulose or inulin diets and receiving 6 Gy irradiation (IR) or no irradiation (nonIR), (B) Data integration (N-integration) analysis of taxonomic and fermentation acid data showing the characteristic microbiota and metabolite profiles of each diet group.

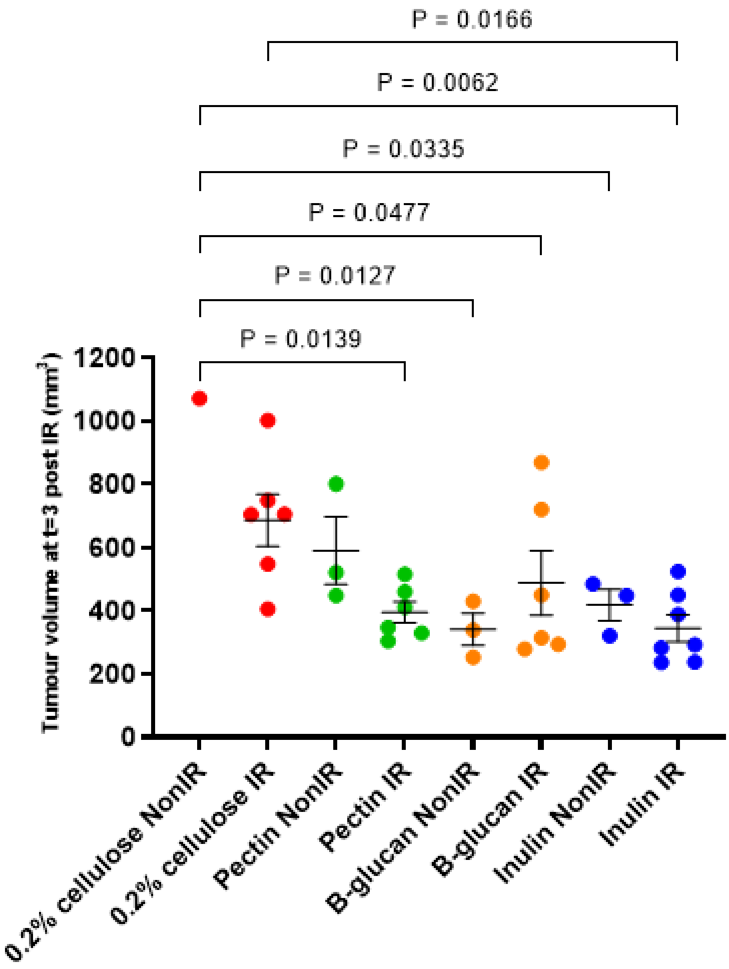

**Figure S4**: **RM-1 tumour growth in C57BL/6J mice fed pectin, β-glucan or inulin.** Comparison of tumour growth at 3 days post-IR (or non-IR equivalent), by one-way ANOVA with Tukey's multiple comparison test. (0.2% cellulose no IR, n=1, IR, n=6; pectin no IR, n=3, IR, n=6; β-glucan no IR, n=3, IR, n=6; inulin no IR, n=3, IR, n=7; missing data due to mice being culled before Day 3 for tumour ulceration +/- rapid growth).

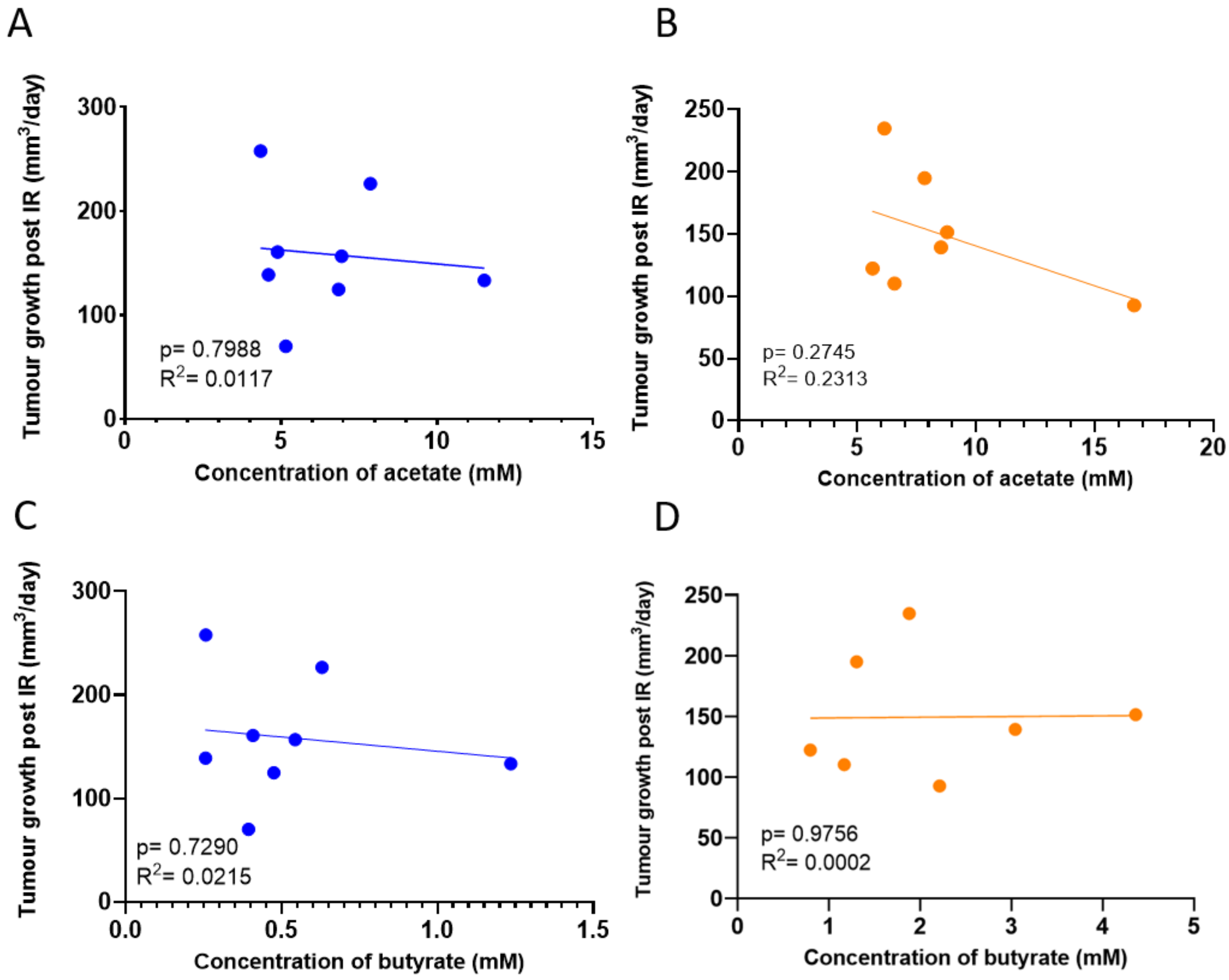

**Figure S5: Correlations of RM-1 tumour growth rates post-IR with acetate and butyrate concentrations post-IR.** Linear regression of: acetate and butyrate concentrations versus rate of tumour growth in inulin- (A) and (C) or B-glucan-fed (B) and (D) post-IR mice by linear regression (inulin: n=8, β-glucan: n=7). Data are presented as mean ± SEM.

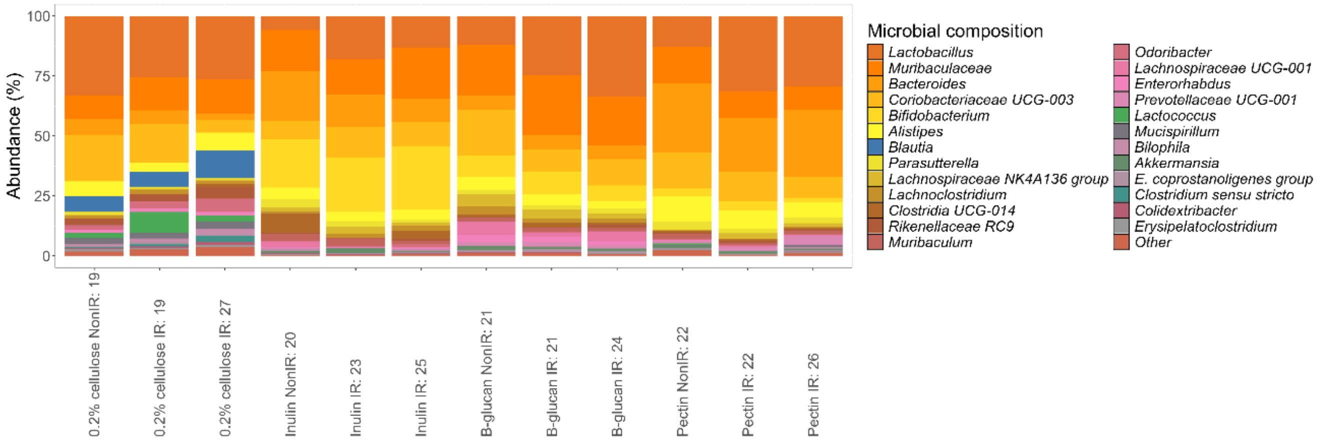

**Figure S6**. Microbiota composition barplot showing the characteristic profiles of each diet group (0.2% cellulose, inulin, B-glucan or pectin diet) considering the effect of cage (cages 19 – 27). The total number of mice per cage were: 19 (n=5), 20 (n=4), 21 (n=5), 22 (n=5), 23 (n=2), 24 (n=5), 25 (n=5), 26 (n=6) and 27 (n=6).

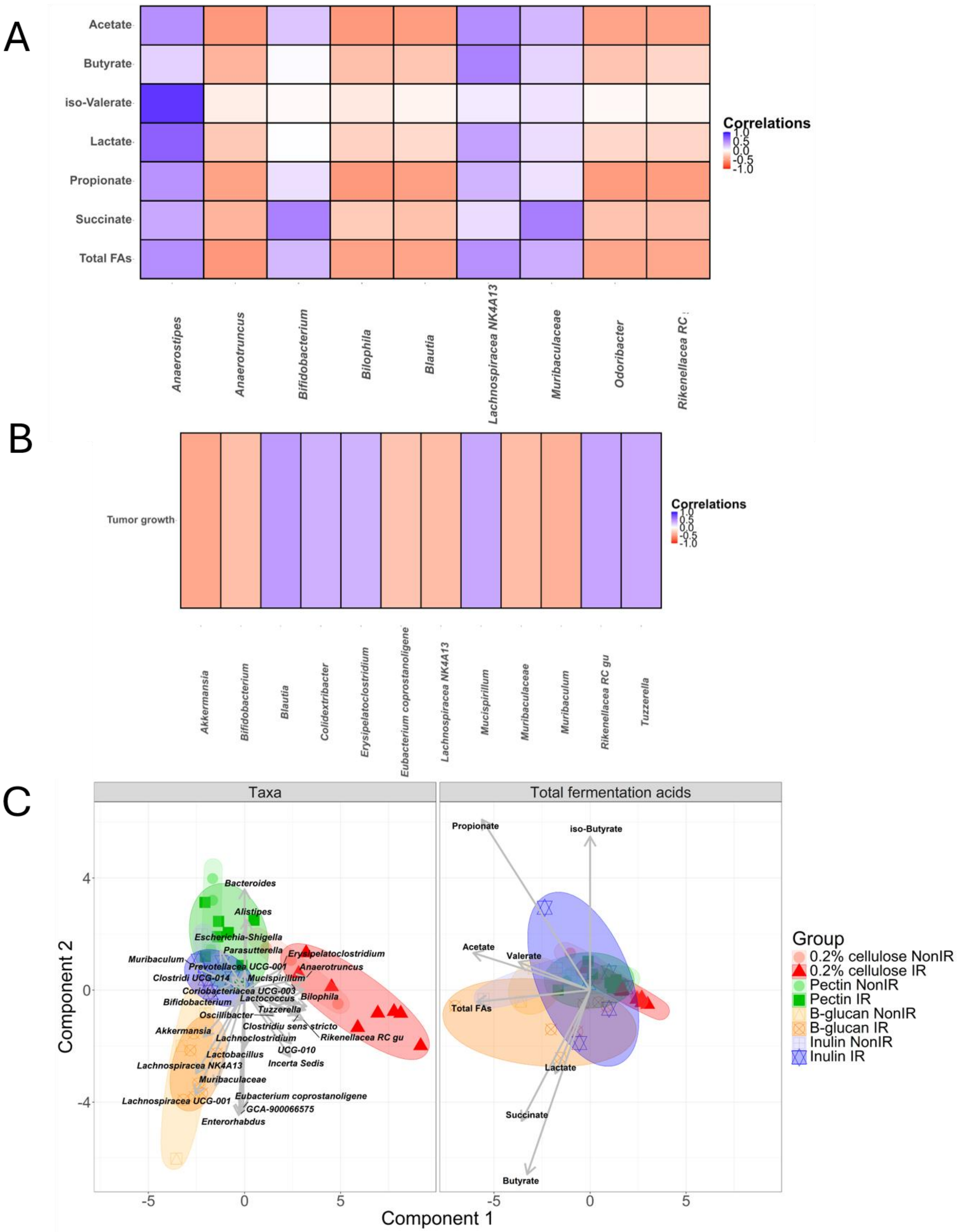
A
Acetate
Butyrate
iso-Valerate
Lactate
Propionate
Succinate
Total FAs
Anaerostipes
Anaerotruncus
Bifidobacterium
Bilophila
Blautia
Lachnospiracea NK4A13
Muribaculaceae
Odoribacter
Rikenellacea RC
Correlations
1.0
0.5
0.0
-0.5
-1.0
B
Tumor growth
Akkermansia
Bifidobacterium
Blautia
Colidextribacter
Erysipelatoclostridium
Eubacterium coprostanoligene
Lachnospiracea NK4A13
Mucispirillum
Muribaculaceae
Muribaculum
Rikenellacea RC gu
Tuzzerella
Correlations
1.0
0.5
0.0
-0.5
-1.0
C
Taxa
Total fermentation acids
Component 2
Component 1
Bacteroides
Alistipes
Escherichia-Shigella
Parasutterella
Erysipelatoclostridium
Muribaculum
Prevotellacea UCG-001
Anaerotruncus
Clostridi UCG-014
Mucispirillum
Coriobacteriacea UCG-003
Bilophila
Bifidobacterium
Lactococcus
Tuzzerella
Oscillibacter
Clostridiu sens stricto
Akkermansia
Lachnoclostridium
Rikenellacea RC gu
UCG-010
Lactobacillus
Incerta Sedis
Lachnospiracea NK4A13
Muribaculaceae
Eubacterium coprostanoligene
Lachnospiracea UCG-001
GCA-900066575
Enterorhabdus
Propionate
iso-Butyrate
Acetate
Valerate
Total FAs
Lactate
Succinate
Butyrate
Group
0.2% cellulose NonIR
0.2% cellulose IR
Pectin NonIR
Pectin IR
B-glucan NonIR
B-glucan IR
Inulin NonIR
Inulin IR

**Figure S7: Correlation heatmaps** showing the associations between relative abundance of specific taxa identified as the dominant/signature taxon in one of the dietary groups (See panel C and Table S3) and (A) fermentation acids or (B) tumour growth rates in tumour-bearing C57BL/6J mice fed 0.2% cellulose, inulin, β-glucan or pectin diets and receiving 6 Gy irradiation (IR) or no irradiation (nonIR), (C) Data integration (N-integration) analysis of taxonomic and fermentation acid data showing the characteristic microbiota and metabolite profiles of each diet group.

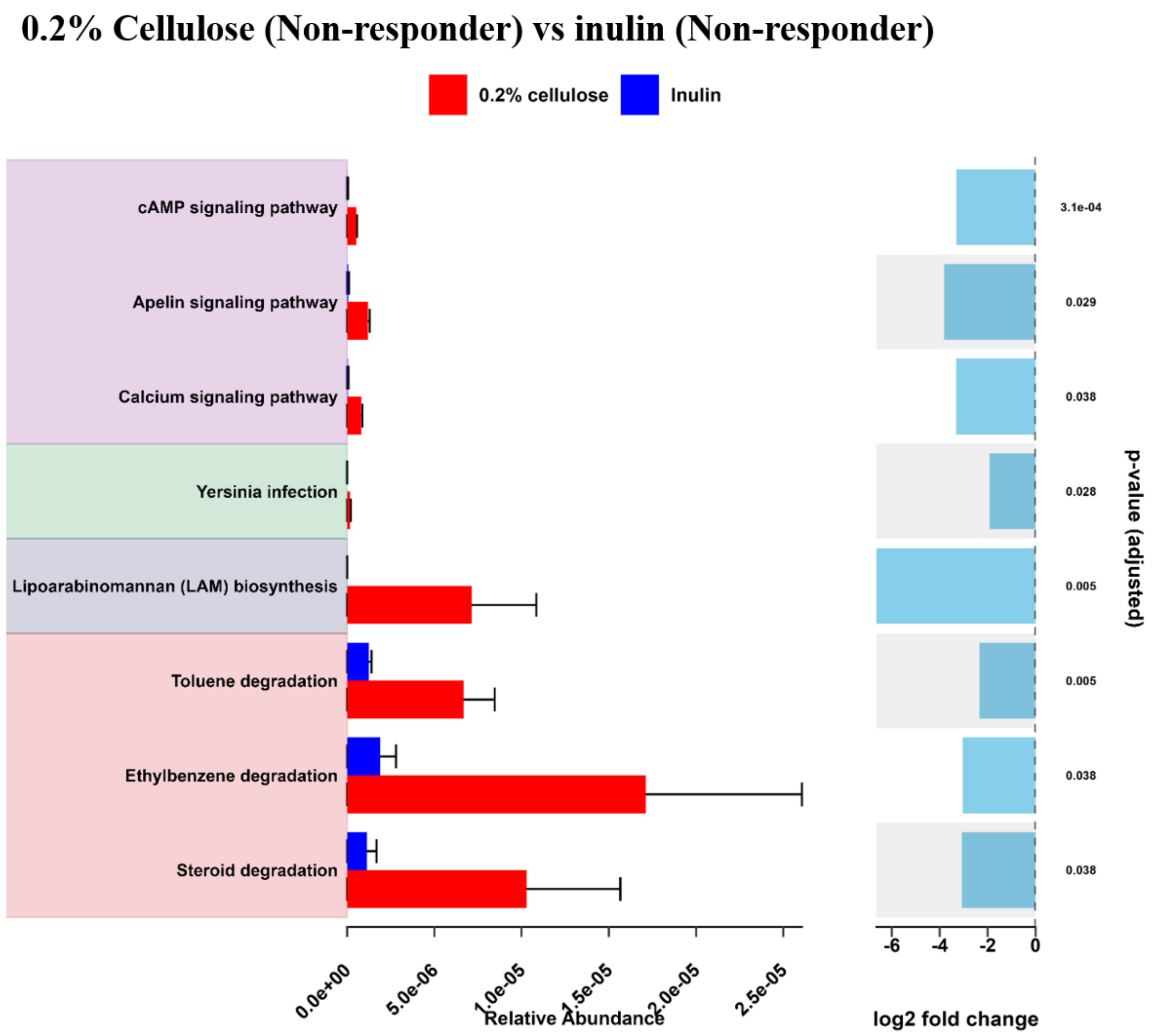

**Figure S8. Predicted microbial metabolic pathways using PICRUSt v.2.0 in non-responder Myc-CaP tumour-bearing FVB mice,** showing statistically significant differences ($p$<0.05) between 0.2% cellulose and inulin diet. These differences were determined using the LinDA (linear model for differential abundance analysis of microbiome compositional data) method.

#### β-glucan (Responder) vs pectin (Responder)

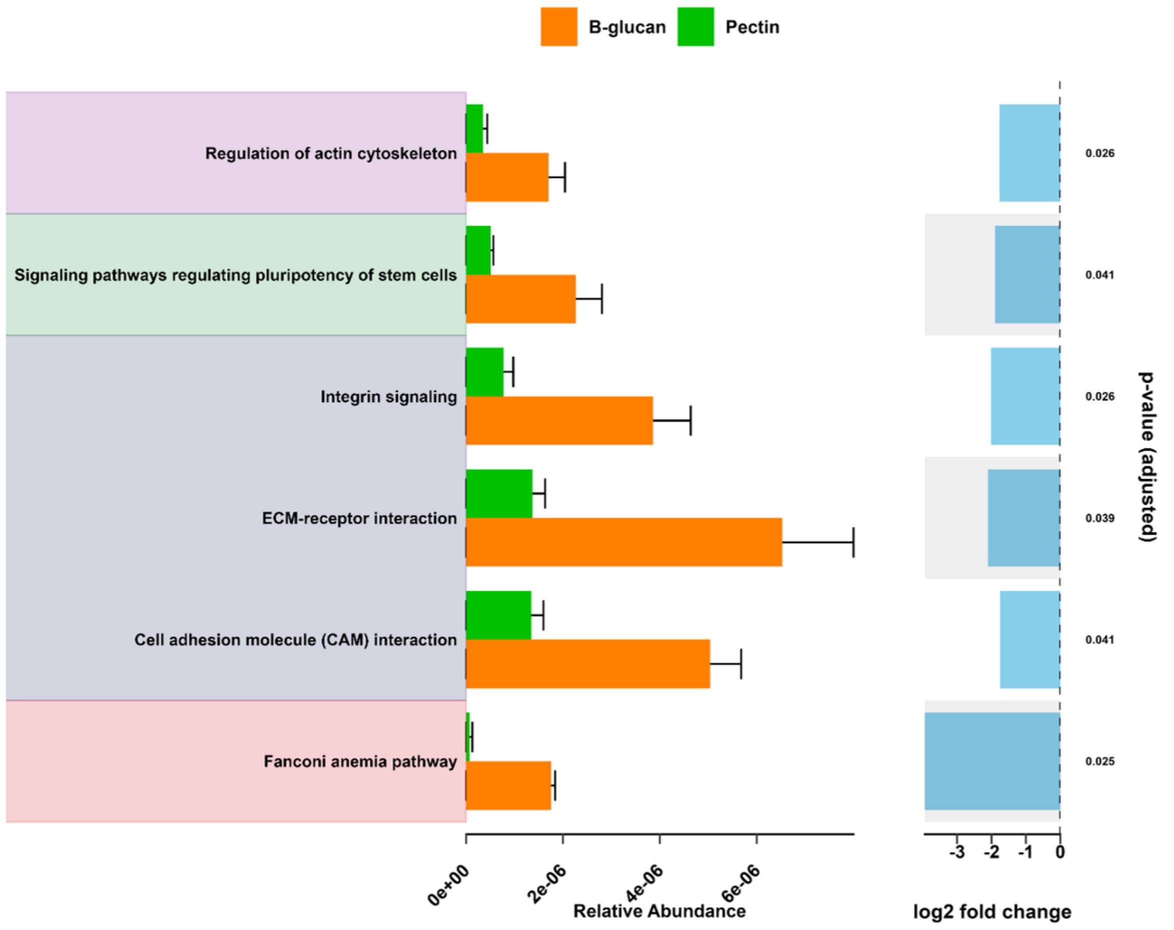

**Figure S9. Predicted microbial metabolic pathways using PICRUSt v.2.0** that show statistically significant differences ($p < 0.05$) between β-glucan and pectin diet groups in responder C57BL/6J mice. These differences were determined using LinDA (linear model for differential abundance analysis of microbiome compositional data) method.

#### Pectin (Non-responder) vs inulin (Non-responder)

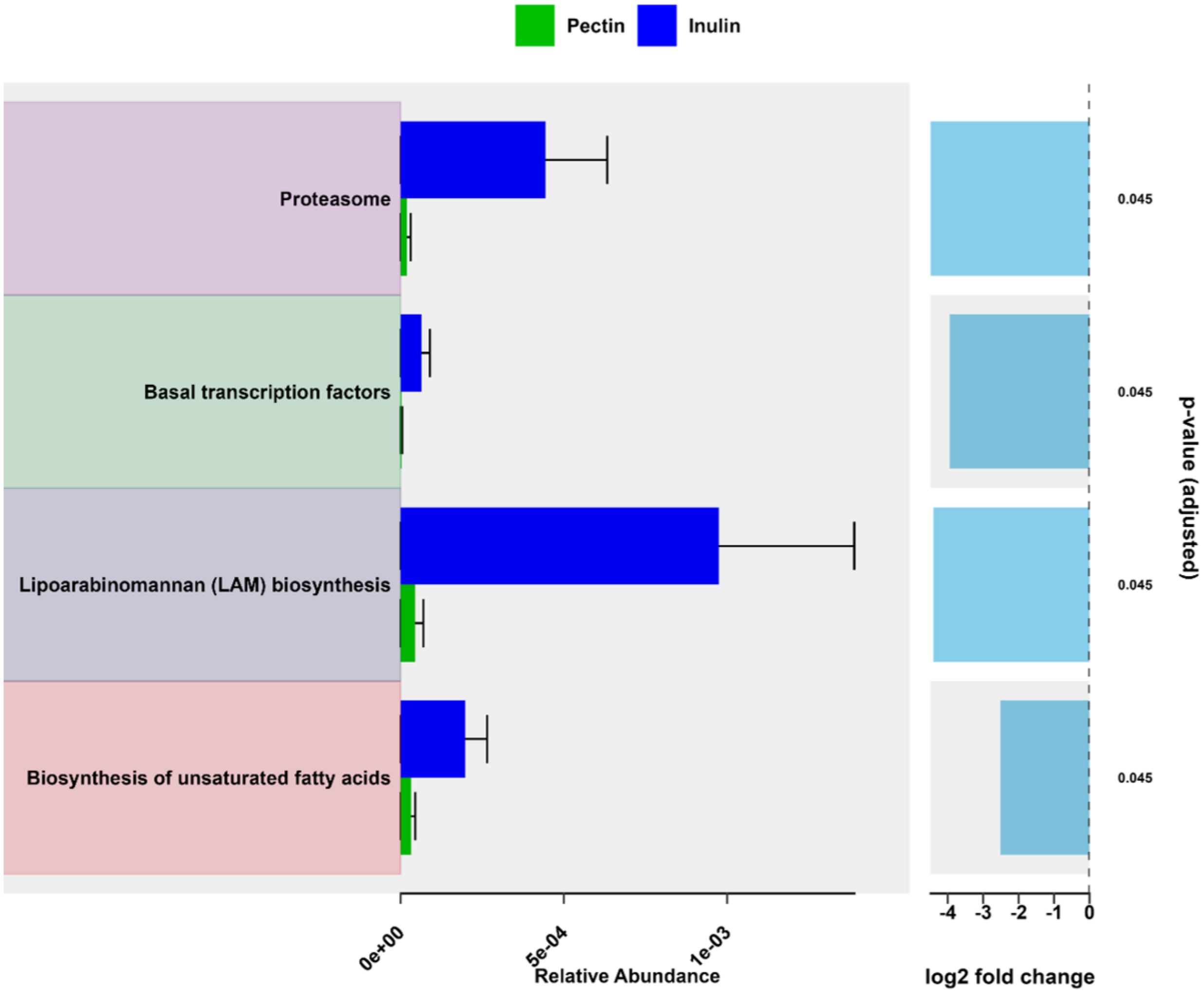

**Figure S10**. Predicted microbial metabolic pathways using PICRUSt v.2.0 that show statistically significant differences ($p<0.05$) between pectin and inulin diet groups in non-responder C57BL/6J mice. These differences were determined using LinDA (linear model for differential abundance analysis of microbiome compositional data) method.

#### β-glucan (Non-responder) vs pectin (Non-responder)

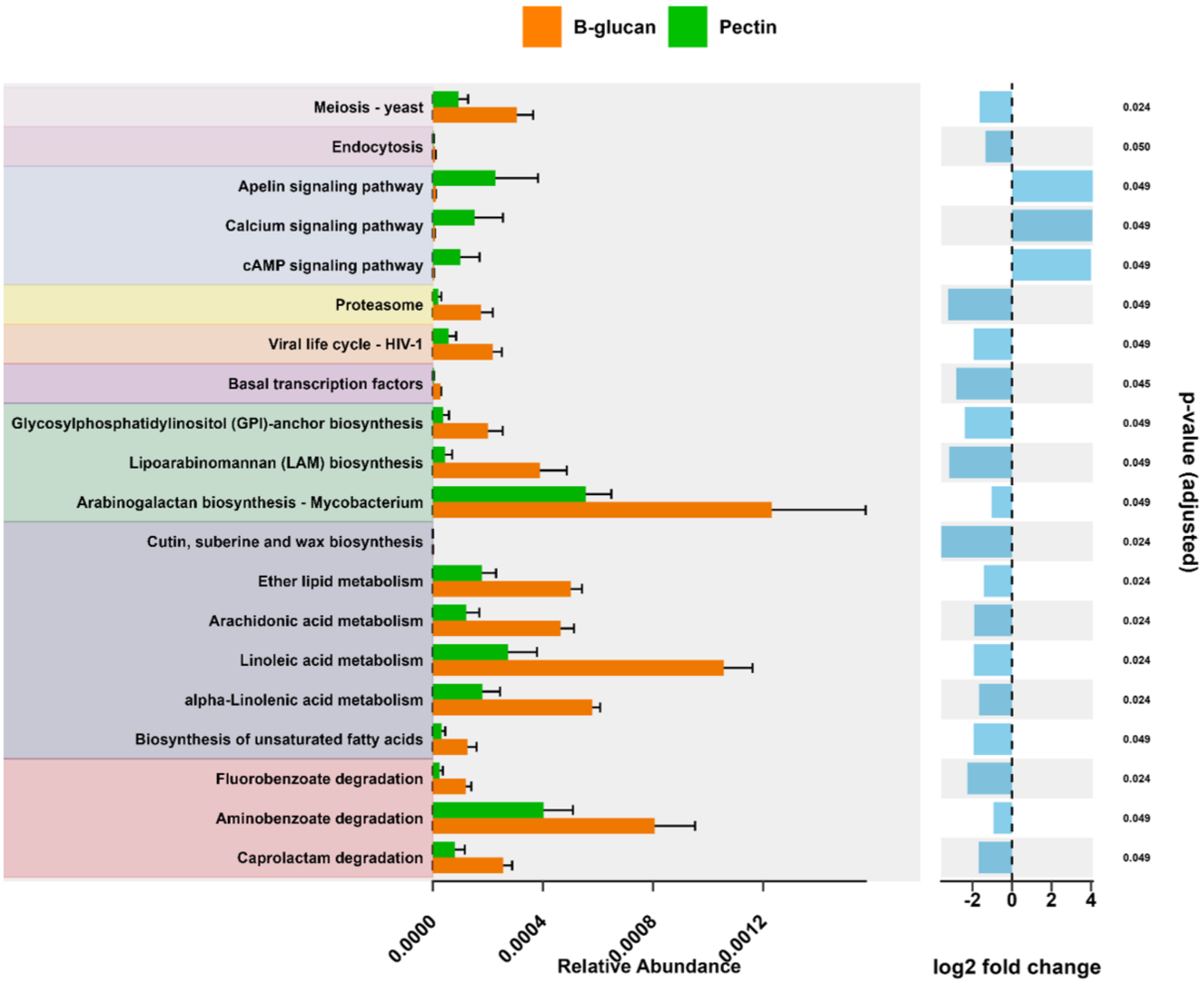

**Figure S11. Predicted microbial metabolic pathways using PICRUSt v.2.**0 that show statistically significant differences ($p < 0.05$) between β-glucan and pectin diet groups in non-responder C57BL/6J mice. These differences were determined using LinDA (linear model for differential abundance analysis of microbiome compositional data) method.

**Table S1**: Formulae for rodent diets used in the study with varying quantities of cellulose, apple pectin, inulin, or β-glucan per-4000 kcal.

| Product | *2gm Cellulose/4000kcal* | *100 gm Inulin/4000kcal* | *101.7 gm Pectin/ 4000 kcal Diet* | *150 gm Orafti bfit/4000kcal* |
|---|---|---|---|---|
| Ingredient | gm | gm | gm | gm |
| Casein | 200 | 200 | 200 | 135 |
| L-Cystine | 3 | 3 | 3 | 3 |
| Corn Starch | 0 | 0 | 0 | 0 |
| Dextrose, Monohydrate | 397.486 | 360 | 346.65 | 285 |
| Maltodextrin 10 | 132 | 132 | 132 | 132 |
| Sucrose | 100 | 100 | 100 | 100 |
| Cellulose, BW200 | 2 | 0 | 0 | 0 |
| Inulin (96%) | 0 | 100 | 0 | 0 |
| Apple Pectin (98.5%) | 0 | 0 | 101.7 | 0 |
| Orafti b-fit flour (22% beta-glucans) | 0 | 0 | 0 | 375 |
| Soybean Oil | 70 | 70 | 70 | 46 |
| tBHQ | 0.014 | 0.014 | 0.014 | 0.014 |
| Mineral Mix S10022G | 35 | 35 | 35 | 35 |
| Vitamin Mix V10037 | 10 | 10 | 10 | 10 |
| Choline Bitartrate | 2.5 | 2.5 | 2.5 | 2.5 |
| FD&C Red Dye #5 | 0 | 0 | 0 | 0 |
| FD&C Yellow Dye #40 | 0 | 0 | 0 | 0 |
| FD&C Blue Dye #1 | 0 | 0 | 0 | 0 |
| Total | 952 | 1012.514 | 1000.864 | 1123.514 |
| Cellulose (gm/~4000 kcal) | 2.0 | 0.0 | 0 | 0.0 |
| Inulin (gm/~4000 kcal) | 0.0 | 96.0 | 0 | 0.0 |
| Pectin (gm/~4000 kcal) | 0.0 | 0.0 | 100.1 | 0.0 |
| Beta-glucans (gm/~4000 kcal) | 0.0 | 0.0 | 0 | 82.5 |
| g | | | | |
| Protein | 177.0 | 177.0 | 177.0 | 176.7 |
| Carbohydrate | 629.5 | 629.5 | 629.5 | 629.5 |
| Fat | 70.0 | 70.0 | 70.0 | 70.0 |
| Total Fibre | 2.0 | 96.0 | 98.5 | 150.0 |
| Protein | *708* | *708* | *708* | *707* |
| Carbohydrate | *2518* | *2518* | *2518* | *2518* |
| Fat | *630* | *630* | *630* | *630* |
| Total (kcal) | *3856* | *3856* | *3856* | *3855* |

**Table S2**. Microbial genera showing high abundance in the taxonomic profiles and identified as microbial biomarkers of MyC-CaP tumour-bearing FVB mice (A) fed 0.2% cellulose or inulin diets ~~(top)~~ or (B) fed 0.2% cellulose or inulin diets and receiving 6 Gy irradiation (IR) or no irradiation (nonIR). Data are expressed as sequence abundance percentage (%). SD: standard deviation. Potential microbial biomarkers ($p < 0.05$ and $p_{adj} < 0.05$) of each group determined by microbiomeMarker statistical methods are highlighted in bold.

**A. MyC-CaP tumour-bearing FVB mice (differences based on diet)**

| **Highest in:** | **Taxa** | **0.2% cellulose (n=7) (Mean)** | **0.2% cellulose (n=7) (SD)** | **Inulin (n=11) (Mean)** | **Inulin (n=11) (SD)** |
|---|---|---|---|---|---|
| 0.2% cellulose | *Blautia* | **8.84** | 2.78 | 0.18 | 0.37 |
| Inulin | Uncultured genus | 3.62 | 1.76 | **10.32** | 8.49 |
| Inulin | *Faecalitalea* | 3.80 | 2.29 | **9.62** | 6.39 |
| Inulin | *Anaerostipes* | 0.36 | 0.12 | **9.53** | 7.74 |

**B. MyC-CaP tumour-bearing FVB mice**

| **Highest in:** | **Taxa** | **Cellulose nonIR (n=3) (Mean)** | **Cellulose nonIR (n=3) (SD)** | **Cellulose IR (n=4) (Mean)** | **Cellulose IR (n=4) (SD)** | **Inulin nonIR (n=3) (Mean)** | **Inulin nonIR (n=3) (SD)** | **Inulin IR (n=8) (Mean)** | **Inulin IR (n=8) (SD)** |
|---|---|---|---|---|---|---|---|---|---|
| Cellulose nonIR | *Parabacteroides* | **61.15** | 2.37 | 53.64 | 5.04 | 46.46 | 3.70 | 53.18 | 12.34 |
| Inulin nonIR | *Faecalitalea* | 3.66 | 0.82 | 3.90 | 3.17 | **14.71** | 8.08 | 7.72 | 4.95 |
| Inulin nonIR | *Anaerostipes* | 0.37 | 0.16 | 0.36 | 0.12 | **10.34** | 7.40 | 9.23 | 8.34 |
| Inulin IR | Uncultured | 2.78 | 1.45 | 4.24 | 1.89 | 5.50 | 4.57 | **12.13** | 9.13 |

**Table S3**. Microbial genera showing high abundance in the taxonomic profiles and identified as microbial biomarkers of RM-1 tumour-bearing C57BL/6J mice (A) fed 0.2% cellulose, inulin, B-glucan or pectin diets, or (B) fed 0.2% cellulose, inulin, B-glucan or pectin diets and receiving 6 Gy irradiation (IR) or no irradiation (nonIR). Data are expressed as sequence abundance percentage (%). SD: standard deviation. ***L.* UCG-001**: *Lachnospiraceae* UCG-001, ***P.* UCG-001**: *Prevotellaceae* UCG-001, **UCG-003**: *Coriobacteriaceae* UCG-003, **UCG-014**: *Clostridia* UCG-014. Potential microbial biomarkers ($p < 0.05$ and $p_{adj} < 0.05$) of each group determined by microbiomeMarker statistical methods are highlighted in bold.

**A. C57BL/6J mice (differences based on diet)**

| **Highest in:** | **Taxa** | **0.2% cellulose (n=11) (Mean)** | **0.2% cellulose (n=11) (SD)** | **Inulin (n=11) (Mean)** | **Inulin (n=11) (SD)** | **Pectin (n=11) (Mean)** | **Pectin (n=11) (SD)** | **β-glucan (n=10) (Mean)** | **β-glucan (n=10) (SD)** |
|---|---|---|---|---|---|---|---|---|---|
| 0.2% cellulose | *Blautia* | **6.70** | 3.11 | 0.00 | 0.00 | 0.00 | 0.00 | 0.04 | 0.10 |
| Inulin | *Bifidobacterium* | 0.03 | 0.06 | **17.01** | 5.47 | 1.99 | 1.23 | 6.92 | 2.39 |
| Inulin | Uncultured genus | 2.89 | 2.14 | **10.63** | 6.40 | 6.97 | 4.98 | 5.62 | 3.83 |
| Pectin | *Bacteroides* | 3.13 | 1.72 | 10.09 | 3.29 | **21.35** | 10.09 | 5.21 | 1.73 |
| Pectin | *Alistipes* | 4.71 | 2.31 | 3.13 | 0.71 | **5.99** | 2.18 | 3.65 | 1.17 |
| B-glucan | *Lactobacillus* | 20.49 | 8.14 | 8.59 | 5.46 | 19.99 | 9.14 | **22.93** | 12.82 |
| B-glucan | *Muribaculaceae* genus | 9.23 | 2.14 | 13.64 | 4.61 | 8.79 | 2.38 | **18.90** | 3.20 |

**B. C57BL/6J mice**

| **Highest in:** | **Taxa** | **Cellulose nonIR (n=3) (Mean)** | **Cellulose nonIR (n=3) (SD)** | **Cellulose IR (n=8) (Mean)** | **Cellulose IR (n=8) (SD)** | **Inulin nonIR (n=3) (Mean)** | **Inulin nonIR (n=3) (SD)** | **Inulin IR (n=8) (Mean)** | **Inulin IR (n=8) (SD)** | **Pectin nonIR (n=3) (Mean)** | **Pectin nonIR (n=3) (SD)** | **Pectin IR (n=8) (Mean)** | **Pectin IR (n=8) (SD)** | **β-glucan nonIR (n=3) (Mean)** | **β-glucan nonIR (n=3) (SD)** | **β-glucan IR (n=7) (Mean)** | **β-glucan IR (n=7) (SD)** |
|---|---|---|---|---|---|---|---|---|---|---|---|---|---|---|---|---|---|
| Inulin NonIR | *Clostridia* UCG-014 | 0.11 | 0.10 | 0.65 | 0.40 | **5.08** | 1.29 | 1.63 | 2.04 | 0.37 | 0.64 | 0.47 | 0.29 | 0.88 | 0.53 | 1.02 | 0.58 |
| Inulin NonIR | *Muribaculum* | 0.04 | 0.08 | 0.00 | 0.00 | **1.90** | 0.11 | 1.32 | 0.93 | 1.51 | 0.11 | 1.14 | 0.66 | 1.55 | 0.65 | 1.48 | 0.29 |
| Inulin IR | *Bifidobacterium* | 0.10 | 0.09 | 0.00 | 0.00 | 12.59 | 0.48 | **18.66** | 5.58 | 2.21 | 1.51 | 1.91 | 1.22 | 7.75 | 1.81 | 6.57 | 2.65 |
| Pectin NonIR | *Parasutterella* | 0.85 | 0.58 | 0.46 | 0.28 | 2.12 | 0.59 | 1.09 | 0.61 | **2.42** | 0.78 | 1.77 | 0.96 | 1.45 | 0.98 | 1.65 | 0.69 |
| Pectin IR | *Bacteroides* | 4.56 | 1.01 | 2.59 | 1.64 | 12.96 | 1.37 | 9.02 | 3.17 | 20.48 | 14.17 | **21.67** | 9.36 | 5.22 | 1.13 | 5.21 | 2.01 |
| Pectin IR | *Prevotellaceae* UCG-001 | 0.00 | 0.00 | 0.02 | 0.07 | 0.00 | 0.00 | 0.01 | 0.04 | 0.85 | 0.58 | **2.62** | 1.26 | 1.32 | 0.13 | 1.19 | 0.54 |
| B-glucan NonIR | *Coriobacteriaceae* UCG-003 | 13.03 | 4.58 | 5.40 | 3.32 | 4.68 | 0.73 | 7.95 | 4.24 | 10.40 | 2.52 | 7.61 | 3.65 | **16.21** | 4.61 | 9.33 | 2.09 |
| B-glucan NonIR | *Lachnospiraceae* UCG-001 | 0.00 | 0.00 | 0.00 | 0.00 | 1.59 | 0.82 | 0.03 | 0.09 | 0.00 | 0.00 | 0.28 | 0.32 | **4.88** | 1.71 | 2.93 | 1.62 |
| B-glucan NonIR | *Lachnoclostridium* | 0.75 | 0.44 | 1.02 | 0.52 | 0.55 | 0.30 | 2.03 | 1.49 | 0.24 | 0.14 | 0.40 | 0.34 | **2.95** | 1.86 | 1.48 | 1.15 |
| B-glucan IR | *Lactobacillus* | 23.32 | 7.78 | 19.43 | 8.53 | 3.63 | 0.89 | 10.46 | 5.27 | 8.89 | 5.13 | 24.15 | 6.27 | 10.62 | 5.14 | **28.20** | 11.38 |
| B-glucan IR | *Muribaculaceae* | 6.71 | 1.86 | 10.17 | 1.36 | 10.89 | 3.29 | 14.67 | 4.77 | 10.53 | 3.90 | 8.14 | 1.39 | 18.25 | 2.85 | **19.18** | 3.51 |

**Table S4. Responder vs non-responder status by cage**. Percentages represent percentages of mice for each individual cage.

| **Cage** | **Diet** | **0.2% cellulose** | |
|---|---|---|---|
| | | **R (%)** | **NR (%)** |
| Cage 19 IR | 0.2% cellulose | 0 (0) | 8 (100%) |
| Cage 27 IR | 0.2% cellulose | 0 (0) | 8 (100%) |
| Cage 20 IR | Inulin | 1 (100%) | 0 (0%) |
| Cage 23 IR | Inulin | 2 (67%) | 1 (33%) |
| Cage 25 IR | Inulin | 2 (50%) | 2 (50%) |
| Cage 21 IR | β-glucan | 2 (100%) | 0 (0%) |
| Cage 24 IR | β-glucan | 1 (25%) | 3 (75%) |
| Cage 22 IR | Pectin | 1 (50%) | 1 (50%) |
| Cage 26 IR | Pectin | 2 (40%) | 3 (60%) |
